# Targeting Noncanonical TEAD Axis to Overcome DNA Repair-Driven Chemoresistance

**DOI:** 10.64898/2026.04.09.717588

**Authors:** Dong-Hyeon Kim, Jongwan Kim, Sangwoo Kang, Hwa-Ryeon Kim, Seok Chan Cho, Haiyan Jin, Inyoung Kim, Eunyoung Jung, Hye-Ran Seo, Eun A Lee, Jung-Soon Mo, Ho Chul Kang, Kyungjae Myung, Jongbum Kwon, Yonghwan Kim, Wantae Kim, Jae-Seok Roe, Donghyuk Shin, Kyoung Tai No, Hyun Woo Park

**Author notes:** Correspondence: Hyun Woo Park, Kyoung Tai No. Equal contributions (Dong-Hyeon Kim, Jongwan Kim).

## Abstract

How transcription factors contribute to DNA damage repair (DDR) independent of canonical gene regulation remains poorly understood. Here, we show that DNA double-strand break (DSB) triggers a functional state transition in TEAD, disengaging it from YAP-dependent transcription and translocating it to DNA lesions as a chromatin-associated DDR factor. Upon genotoxic stress, TEAD is rapidly recruited to damaged chromatin through a biphasic mechanism involving early poly(ADP-ribose)-dependent recruitment followed by ATM-driven γH2AX-mediated retention. At DNA lesions, TEAD constrains chromatin over-relaxation, suppresses excessive end resection, and promotes non-homologous end joining (NHEJ) independent of its canonical function. Conserved residues within the TEA domain mediate this noncanonical chromatin engagement of TEAD and define a druggable N-terminal interface. Pharmacological targeting of this interface impaired TEAD-dependent DSB repair and sensitized tumors to chemotherapy. Together, these findings establish noncanonical TEAD as a critical DNA repair factor and a clinically actionable driver of chemoresistance, while providing a conceptual framework for understanding how broader families of transcription factors may be repurposed as targetable DNA damage regulators in cancer.

## Introduction

Maintenance of genome integrity requires coordinated DNA damage signaling and repair within a dynamically regulated chromatin environment^1, 2^. Following DNA double-strand breaks (DSBs), chromatin undergoes rapid and highly orchestrated structural transitions characterized by transient relaxation and subsequent re-compaction, that determine repair factor accessibility, pathway choice, and cell fate^3, 4^. Although the enzymatic machinery of the DNA damage response (DDR)^5–7^ and the roles of canonical chromatin remodelers have been extensively characterized^3, 8–10^, it remains unclear whether sequence-specific DNA-binding proteins such as transcription factors can be directly repurposed at sites of DNA damage to control chromatin repair. This question is particularly important in cancer, where transcription factors are frequently dysregulated and genotoxic stress is a defining selective pressure imposed by both tumor evolution and therapy.

Accumulating evidence suggests that transcription factors can be localize to damaged chromatin and influence repair independently of their transcriptional output^11–15^. However, the mechanistic basis, physiological significance, and therapeutic relevance of such noncanonical functions remain poorly defined. In particular, it is unknown whether oncogenic transcription factors can undergo regulated functional transitions in response to DNA damage and whether these transitions support tumor survival under genotoxic stress. Resolving this issue would not only broaden our understanding of genome maintenance but also identify a class of cancer vulnerabilities driven by the non-transcriptional functions of transcription factors.

The TEA domain transcription factors, TEAD1-4, provide a compelling candidate in which to address this question^16, 17^. TEAD proteins are classically viewed as transcriptional effectors of the Hippo pathway that cooperate with YAP and TAZ and bind M-CAT motifs in the target gene promotors to drive proliferative and tumor-promoting gene expression programs^18–20^. Accordingly, TEAD-mediated transcription is most evident in tumors harboring Hippo pathway alterations that induce YAP/TAZ hyper-activation, therefore therapeutic efforts are largely being focused on blocking the canonical YAP/TAZ-TEAD signaling^21–23^. Yet this view may be incomplete, because TEAD proteins remain broadly expressed across cancers, including contexts in which canonical Hippo pathway output and YAP/TAZ expression are weak or dispensable, raising the possibility that TEAD may have physiologically important functions outside its established transcriptional role^21, 24–30^. Whether TEAD can be repurposed at sites of DNA damage as a direct chromatin-associated regulator, and whether such functions contribute to chemoresistance has not been determined.

Here, we show that DNA double-strand break triggers a functional state transition in TEAD, disengaging it from canonical transcriptional activity and repurposing it as a chromatin-associated regulator of DNA repair. We find that TEAD is recruited to damaged chromatin through sequential poly(ADP-ribose)-dependent recruitment and ATM-driven γH2AX-mediated retention, where it restrains chromatin over-relaxation, suppresses excessive end resection, and promotes non-homologous end joining. These findings uncover a previously unrecognized pathophysiological role for the noncanonical TEAD in maintaining genome integrity during genotoxic stress and identify a clinically relevant mechanism underlying chemoresistance that is distinct from the canonical Hippo signaling.

This work provides a rationale for developing a new class of N-terminal TEAD inhibitor that targets the noncanonical DNA repair function rather than its conventional transcriptional activity, thereby expanding the therapeutic scope of TEAD-directed treatment strategies to overcome chemotherapy resistance. More broadly, our study establishes TEAD as a proof-of-principle transcription factor that can be repurposed as direct chromatin regulators at DNA lesions, highlighting non-transcriptional DNA repair functions as a potentially targetable driver of cancer vulnerability.

## Results

### Canonical chromatin engagement of TEAD requires direct YAP/TAZ interaction

Although transcription factors are well-established modulators of chromatin architecture during transcription^31, 32^, whether they can repurpose this chromatin-engaging capacity to translocate to DNA lesions and orchestrate DNA repair independently of gene regulation remains largely unexplored. TEAD provides a useful system for this purpose because their canonical chromatin association to M-CAT motifs at target gene promotors is tightly coupled to YAP and TAZ co-activators, allowing chromatin engagement to be experimentally separated from protein abundance and other indirect transcriptional outputs^33, 34^.

We first tested whether canonical TEAD chromatin association is indeed dependent on YAP/TAZ. Depletion of YAP/TAZ did not alter total nuclear TEAD signal under standard fixation conditions, indicating that YAP/TAZ loss does not affect TEAD protein abundance. However, when cells were subjected to CSK pre-extraction before fixation to enrich for chromatin-retained nuclear factors, the residual pan-TEAD signal was markedly diminished by YAP/TAZ depletion, indicating TEAD is not stably retained on chromatin in the absence of these co-activators (**Figure 1A and 1B**). Consistently, genome-wide TEAD4 ChIP-seq analysis in wild-type and YAP/TAZ-knockout cells further showed selective loss of TEAD4 occupancy at promoters and enhancers of canonical Hippo target genes, whereas no evident gain of TEAD4 binding was detected at alternative genomic regions (**Figure 1C, Figure S1A, and S1B**). This dependency was also observed under physiological and pharmacologic conditions that suppress YAP/TAZ activity^29^, including serum starvation^35^, cytoskeletal disruption with latrunculin B^20, 36^, and Src inhibition with dasatinib^37^, which each reduced the chromatin-bound TEAD pool (**Figure S1C and S1D**). In parallel, non-extracted samples confirmed a corresponding reduction in nuclear YAP/TAZ signaling states (**Figure S1E and S1F**). These findings further support the conclusion that basal TEAD chromatin engagement is tightly coupled to nuclear YAP/TAZ availability.

**Figure 1.**
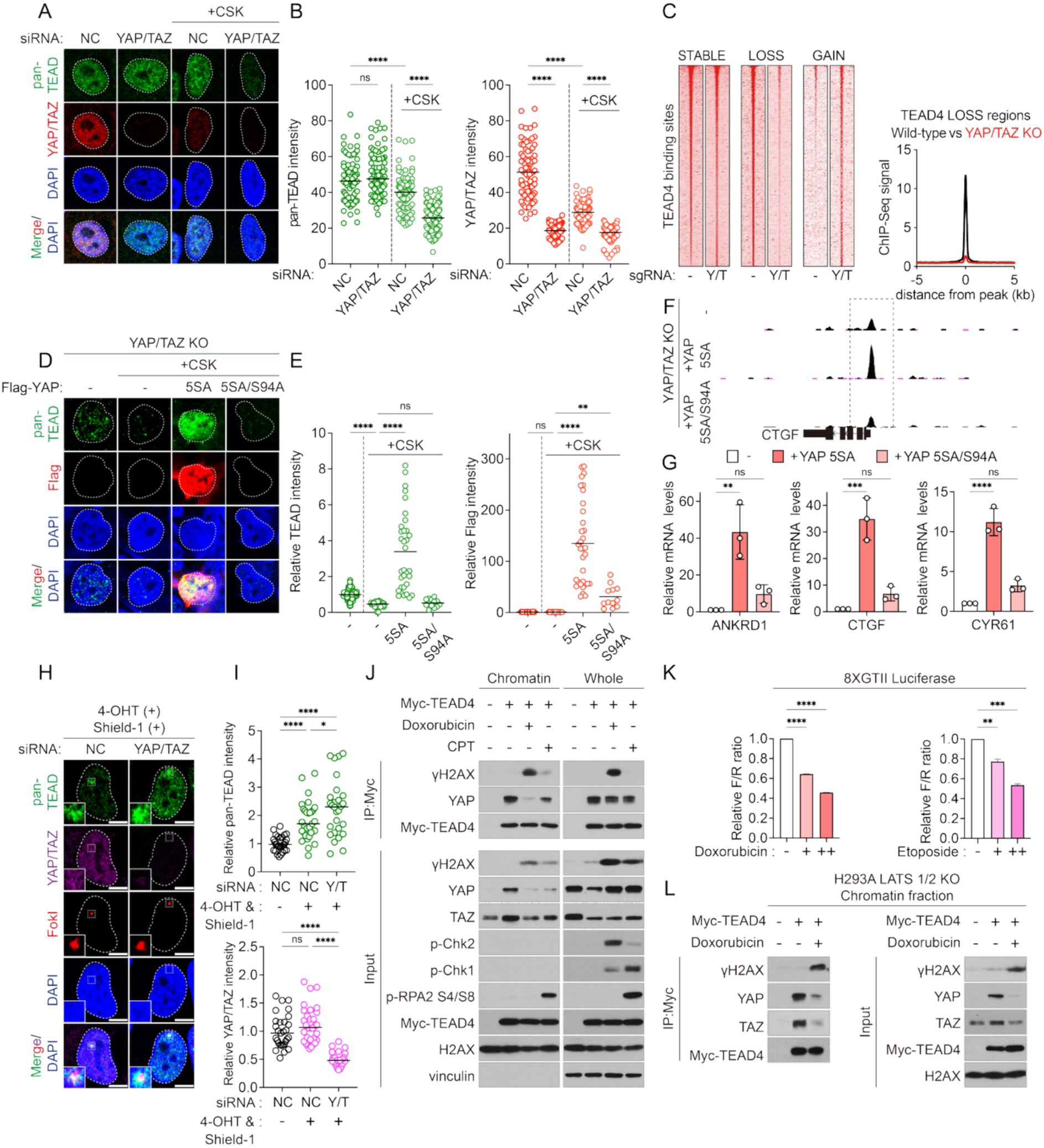
DNA double-strand breaks induce a noncanonical chromatin state of TEAD. (A) Confocal images of endogenous pan-TEAD (green) and YAP/TAZ (red) in U2OS cells transfected with non-targeting control siRNA (siNC, 20 nM) or combined YAP/TAZ siRNA (siYAP 10 nM + siTAZ 10 nM). “+CSK” indicates samples pre-extracted with CSK buffer prior to fixation to enrich chromatin-associated proteins. (B) Quantification of nuclear pan-TEAD and YAP/TAZ fluorescence intensities from (A). Lines indicate medians (siNC: n = 80; siYAP/TAZ: n = 88; +CSK siNC: n = 100; +CSK siYAP/TAZ: n = 105). (C) TEAD4 ChIP-seq profiles comparing wild-type versus YAP/TAZ-knockout (KO) HEK293A cells. Left: Genomic tracks highlighting regions with stable, lost, or gained TEAD4 binding. Right: Average ChIP-seq signal density at TEAD-LOSS regions. (D) Confocal images of HEK293A YAP/TAZ-knockout cells reconstituted with empty vector, FLAG-YAP-5SA, or FLAG-YAP-5SA/S94A. Cells were stained for pan-TEAD (green) and FLAG (red). (E) Quantification of relative nuclear intensities of pan-TEAD and FLAG in cells from (D), normalized to non-extracted (–CSK) controls. Box-and-whisker plots show minimum–maximum ranges (–CSK control: n = 195; +CSK control: n = 58; YAP-5SA: n = 33; YAP-5SA/S94A: n = 13). (F) TEAD4 ChIP-seq signal at the CTGF promoter in YAP/TAZ-knockout HEK293A cells transfected as in (D). Data from two independent experiments. (G) qPCR analysis of canonical Hippo target genes (ANKRD1, CTGF, CYR61) in YAP/TAZ-knockout HEK293A cells expressing empty vector, YAP-5SA, or YAP-5SA/S94A. Expression values were normalized to control (three independent experiments). (H) Recruitment of pan-TEAD and YAP/TAZ to FokI-induced DNA double-strand breaks (DSBs) in U2OS 2-6-5 cells transfected with non-targeting control siRNA (siNC, 20 nM) or combined siYAP (10 nM) and siTAZ (10 nM). (I) Quantification of pan-TEAD and YAP/TAZ fluorescence intensities at FokI-positive regions, normalized to FokI-negative regions. Lines indicate medians (negative, n = 29; siNC, n = 31; siY/T, n = 28). (J) Immunoblot analysis of Myc–TEAD4 immunoprecipitates from chromatin and whole-cell fractions in HEK293A TEAD1/2/4-knockout cells transfected with Myc–TEAD4 and treated with doxorubicin (5 μM, 2 h) or camptothecin (CPT; 1 μM, 2 h). (K) 8×GTIIC luciferase reporter assay measuring YAP/TAZ–TEAD transcriptional activity in HEK293A wild-type cells treated with doxorubicin or etoposide at 1 μM (+) or 3 μM (++) for 8h. Data represent two biological replicates for etoposide and three for doxorubicin. (L) Immunoblot analysis of Myc–TEAD4 immunoprecipitates from chromatin and whole-cell fractions in HEK293A LATS1/2-knockout cells transfected with Myc–TEAD4 and treated with doxorubicin (5 μM, 2 h).

Because YAP has been proposed to stabilize TEAD on chromatin^33^, we tested whether TEAD retention requires physical YAP interaction. In YAP/TAZ-knockout cells, re-expression of constitutively active YAP-5SA restored chromatin-associated pan-TEAD signal after CSK extraction, but not the TEAD-binding-deficient mutant YAP-5SA/S94A (**Figure 1D and 1E**). This requirement for direct interaction was recapitulated at both chromatin occupancy and transcriptional output levels. YAP-5SA restored TEAD4 binding at the CTGF promoter and rescued expression of canonical Hippo target genes, including ANKRD1, CTGF, and CYR61, while YAP-5SA/S94A failed to rescue any of these effects (**Figure 1F and 1G**). Together, these results demonstrate that canonical TEAD chromatin engagement is not intrinsic to TEAD itself, but requires direct YAP/TAZ interaction to sustain its transcription-coupled occupancy at target loci

Importantly, this strict YAP/TAZ dependence establishes a clearly defined canonical TEAD chromatin state that can be experimentally uncoupled. This provided the rationale to test whether DNA damage induces a distinct mode of TEAD chromatin engagement, separate from its established role in YAP/TAZ-dependent transcription.

### DNA double-strand breaks induce a noncanonical chromatin state of TEAD

Having established that canonical TEAD chromatin engagement requires direct YAP/TAZ interaction, we next asked whether DNA damage triggers a distinct mode of TEAD chromatin association. If TEADs were repurposed during genotoxic stress, DNA damage would be expected not only to recruit TEAD to lesions, but also to disengage it from its canonical transcription-associated chromatin state.

To test whether DNA damage triggers functional state transition, we examined TEAD localization following DSB induction. Using U2OS 2-6-5 cells harboring an inducible FokI nuclease system^38^, we observed robust accumulation of TEADs at site specific DSBs, whereas YAP/TAZ showed no detectable recruitment (**Figure 1H and 1I**). Notably, TEAD enrichment at laser-induced damage tracks was further enhanced in YAP/TAZ-deficient cells, indicating that TEAD recruitment to damaged chromatin occurs independently of YAP/TAZ and may even be facilitated when TEAD is released from canonical transcriptional complexes (**Figures S2A and S2B**).

To determine whether TEAD redistribution reflects a chromatin-state transition, we next examined TEAD complex organization after DSB induction. Chromatin fractionation followed by immunoprecipitation showed that DNA damage selectively disrupted TEAD–YAP/TAZ complexes within the chromatin fraction, concomitant with increased TEAD association with γH2AX-marked chromatin, whereas total cellular TEAD–YAP/TAZ interactions remained largely unchanged (**Figure 1J**). Notably, this chromatin-specific redistribution was more pronounced following doxorubicin, which generates blunt-ended DSBs, than by camptothecin (CPT).

Because TEAD release from YAP/TAZ on chromatin should attenuate canonical transcriptional output^33, 34^, we examined TEAD-dependent reporter activity under genotoxic stress. Both doxorubicin and etoposide markedly suppressed 8×GTIIC luciferase activity, indicating that DSB induction is accompanied by functional disengagement of TEAD from YAP/TAZ-driven transcription^20^ (**Figure 1K**). Importantly, disruption of TEAD–YAP/TAZ chromatin association and suppression of TEAD-dependent transcription were also observed in LATS1/2-deficient cells, indicating that this switch occurs independently of canonical Hippo kinase signaling (**Figure 1L and S2C**). Instead, inhibition of upstream DDR kinases, including ATM and PARP^39–41^, restored YAP/TAZ retention on the chromatin during DSB induction, demonstrating that this redistribution is driven by DNA damage signaling rather than by upstream Hippo pathway status (**Figure S2D**).

Together, these findings demonstrate that DNA damage induces a noncanonical chromatin state of TEAD, characterized by its release from YAP/TAZ-dependent transcriptional complexes and redistribution to damaged chromatin. This damage-induced reprogramming establishes TEAD as a candidate chromatin-associated DDR factor and provides the mechanistic basis to investigate how TEAD is recruited to DNA lesions and how it contributes to DNA repair and chemoresistance

### TEAD engages DNA lesions through a biphasic TEA domain-dependent mechanism

The DSB-induced noncanonical chromatin state of TEAD prompted us to ask how TEAD physically associates with DNA lesions. If TEAD functions directly at DNA lesions, we hypothesized that its recruitment would be rapid, conserved across paralogs, and dependent on defined structural features rather than YAP-binding motif. Live-cell imaging revealed that TEAD4 rapidly accumulated at γH2AX-marked damage tracks within minutes and remained enriched for up to 2 hours after irradiation (**Figure S3A**). All four TEAD paralogs showed comparable recruitment kinetics, with detectable localization within approximately 30 seconds, indicating that rapid damage-site engagement is a conserved property of the TEAD family and their recruitment is an early and sustained component of the DNA damage response (**Figure 2A and 2B**).

**Figure 2.**
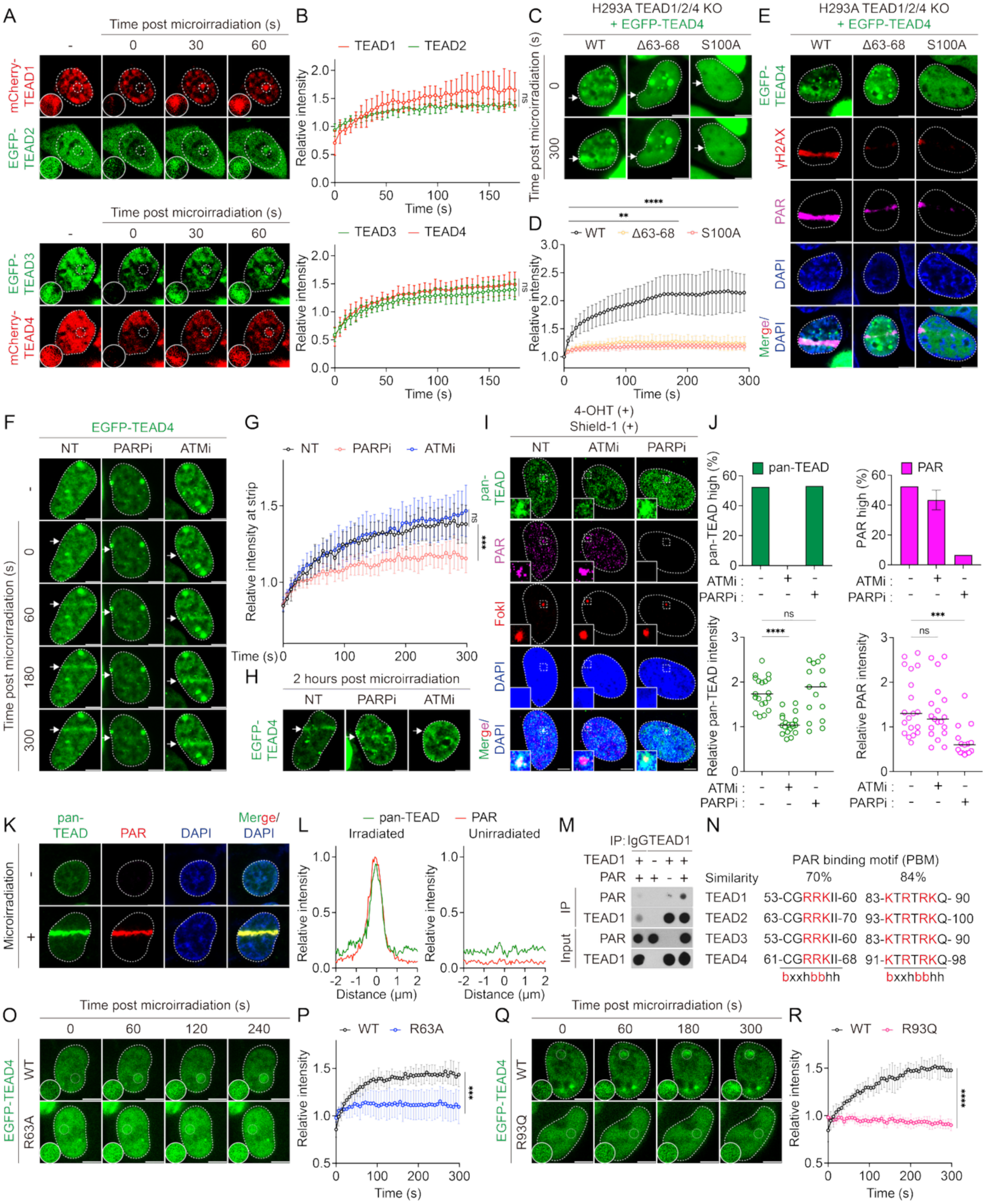
TEA domain directs biphasic recruitment of TEAD to DNA lesions through PARP and ATM signaling. (A) Live-cell confocal imaging showing rapid accumulation of mCherry–TEAD1, EGFP–TEAD2, EGFP–TEAD3, and mCherry–TEAD4 to laser-induced DNA damage sites within 1 min. White dashed circles indicate irradiated regions. (B) Quantification of TEAD paralog recruitment kinetics at microirradiated regions, with fluorescence intensities normalized to the pre-irradiation intensity of the irradiated region (mean ± SD; n = 5 for mCherry–TEAD1 and EGFP–TEAD2; n = 6 for EGFP–TEAD3 and mCherry–TEAD4). (C) Live-cell imaging of EGFP–TEAD4 WT, Δ63–68, or S100A expressed in HEK293A TEAD1/2/4-knockout cells following microirradiation (0 and 300 s). White arrows indicate irradiated regions. (D) Quantification of fluorescence intensity at irradiated regions from (C), normalized to pre-irradiation intensity (WT, n = 5; Δ63–68, n = 7; S100A, n = 4; mean ± SD). (E) Confocal images of EGFP–TEAD4 WT, Δ63–68, or S100A (green) co-stained with γH2AX (red) and PAR (magenta) 2 h after microirradiation. (F) Live-cell imaging of EGFP–TEAD4 expressed in HEK293A TEAD1/2/4-knockout cells pretreated with PARP inhibitor (olaparib, 5 μM) or ATM inhibitor (KU-60019, 5 μM) for 30 min before laser microirradiation. White arrows indicate irradiated regions. (G) Quantification of EGFP–TEAD4 recruitment dynamics at microirradiation tracks, normalized to pre-irradiation intensity (mean ± SD; NT, n = 7; PARPi, n = 8; ATMi, n = 10). (H) Confocal images of EGFP–TEAD4–expressing HEK293A TEAD1/2/4-knockout cells 2 h after microirradiation under the conditions shown in (G). (I) Recruitment of pan-TEAD and poly(ADP-ribose) (PAR) to FokI-induced DSBs in U2OS 2-6-5 cells treated with ATM inhibitor (KU-60019, 5 μM) or PARP inhibitor (olaparib, 5 μM). (J) Fraction of cells showing enrichment of pan-TEAD (green) or PAR (magenta) at FokI-positive regions (two independent biological replicates; upper), and quantification of fluorescence intensities at FokI-positive regions normalized to FokI-negative regions. Lines indicate medians (NT, n = 19; ATMi, n = 19; PARPi, n = 15). (K) Confocal images showing co-localization of endogenous pan-TEAD with poly(ADP-ribose) (PAR) at laser-induced DNA damage tracks. (L) Line-scan profiles showing relative pan-TEAD (green) and PAR (red) intensities across damaged and undamaged regions. (M) In vitro pull-down assay demonstrating binding of recombinant TEAD1 to poly(ADP-ribose) (PAR) chains following TEAD1 immunoprecipitation. (N) In silico motif analysis of the conserved TEA domain in TEAD1–4 using the PROSITE pattern -[CGAVILMFYW]-[FILPV] (PATTINPROT, NPS@), identifying motifs enriched in basic residues (highlighted in red). (O) Live-cell imaging of EGFP–TEAD4 WT versus PAR-binding–deficient mutants R63A expressed in TEAD1/2/4-knockout cells. Dashed circles indicate irradiated regions. (P) Quantification of EGFP–TEAD4 recruitment kinetics for WT versus R63A, normalized to pre-irradiation intensity of the same region (mean ± SD; WT vs. R63A: n = 8 and 10) (Q) Live-cell imaging of EGFP–TEAD4 WT versus PAR-binding–deficient mutants R93Q expressed in TEAD1/2/4-knockout cells. Dashed circles indicate irradiated regions. (R) Quantification of EGFP–TEAD4 recruitment kinetics for WT versus R93Q, normalized to pre-irradiation intensity of the same region (mean ± SD; WT vs. R93Q: n = 6 and 8). Unless otherwise indicated, scale bars = 5 μm. Error bars represent SD. Statistical significance was determined by one-way ANOVA: ns, not significant; *P < 0.05; **P < 0.01; ***P < 0.001; ****P < 0.0001. See also Figure S3.

To test whether noncanonical TEAD localization requires its intrinsic DNA-binding function of the TEA domain^11, 13^, we examined TEAD4 mutants defective in TEA domain-mediated DNA interaction^16^. TEAD4 Δ63–68 (partial L1 loop deletion) and S100A (α3 helix mutant) failed to accumulate at laser-induced damage tracks at both early and late time points (**Figure 2C and 2D**). Consistently, both mutants were defective in localization to microirradiated chromatin marked by PAR and γH2AX at later time points (**Figure 2E**). These results indicate that an intact TEA domain is required for stable association of TEAD with damaged chromatin.

Because poly(ADP-ribose) polymerase (PARP)-dependent PARylation and ATM-dependent γH2AX formation are among the earliest chromatin modifications induced by DSBs^40, 42^, we next asked how these pathways differentially regulate TEAD recruitment. PARP inhibition with olaparib markedly impaired TEAD accumulation during the first 2 min after laser damage, whereas ATM inhibition had little effect during this early phase (**Figures 2F and 2G**). In contrast, sustained TEAD retention at later time points required ATM activity but was largely insensitive to PARP inhibition (**Figure 2H**). This temporal separation suggested that TEAD is recruited to DSBs through mechanistically distinct early and late phases. Consistent results were observed in the FokI-induced site-specific DSB system, where ATM inhibition abolished TEAD retention at induced breaks, whereas PARP inhibition did not prevent late TEAD accumulation despite effective suppression of detectable PAR (**Figures 2I and 2J**). Together, these findings define a biphasic recruitment mechanism in which TEAD is initially recruited through PARP-dependent chromatin modification and subsequently stabilized through ATM-dependent γH2AX signaling. These findings identify the TEA domain as a dual-function chromatin interface that couples rapid damage sensing to sustained DNA lesion retention, thereby enabling TEAD to execute a noncanonical DNA repair program.

### A basic interface in the TEA domain mediates PAR-dependent recruitment of TEAD

Because early recruitment required PARP activity^11, 40^, we tested whether TEAD directly binds PAR chains. Following laser microirradiation, PAR chains were generated within seconds, as indicated by rapid accumulation of the PAR sensor mRuby2-PBZ peaking at approximately 10 seconds^43^, preceding maximal TEAD enrichment at approximately 60 seconds (**Figure S3B and S3C**). Endogenous TEAD selectively co-localized with PAR at laser-induced damage tracks but not in undamaged chromatin (**Figure 2K and 2L**). In line with this observation, recombinant TEAD bound purified PAR chains in vitro (**Figure 2M**), supporting a direct biochemical interaction.

To identify the residues mediating PAR recognition, we performed in silico motif analysis of the TEA domain, focusing on canonical PAR-binding motifs (PBMs) enriched in basic residues^44, 45^. Two candidate PBMs were identified within the TEA domain: CGRRKIIL in the L1 loop and KTRTRKQV in the adjacent α3 helix (**Figure 2N**), consistent with recently reported TEAD4 PBMs^46^. To distinguish PAR recognition from DNA-binding function, we individually mutated each basic residue (R63A, R64A, K65A; R93Q). Among these, R63A partially impaired recruitment, whereas R93Q completely abolished TEAD accumulation at laser-induced damage tracks (**Figure 2O-R**). In contrast, R64A and K65A showed no detectable defect (**Figures S3D and S3E**). These results indicate that conserved basic residues within the TEA domain contribute differentially to PAR-dependent recruitment, with R93 functioning as a dominant determinant of damage-site localization.

Structural modeling using AlphaFold3 positioned these residues within a basic pocket spanning the L1 loop and adjacent α3 helix, consistent with electrostatic engagement of PAR chains (**Figure S3F**). Although the model did not predict direct contact between R93 and PAR, the complete loss of recruitment in the R93Q mutant suggests that R93 plays a structural role in stabilizing the PAR-recognition interface. Together, these data define a PAR- dependent recruitment mechanism mediated by a conserved basic interface within the TEA domain.

### Early TEAD recruitment restrains chromatin relaxation and end resection to promote NHEJ

PARP activation induces transient chromatin relaxation at DNA double-strand breaks (DSBs), necessitating subsequent factors to re-establish local chromatin structure^4, 5, 8, 9^. Because TEAD is rapidly recruited to DSBs through PAR recognition, we asked whether it contributes to early chromatin condensation^10^.

To directly assess chromatin dynamics after DNA damage, we monitored PAGFP–H2B following laser microirradiation^47^. TEAD depletion resulted in immediate and progressively enhanced chromatin expansion at damage tracks (**Figure 3A and 3B**). Re-expression of wild-type TEAD4 restored normal compaction kinetics in TEAD1/2/4-knockout cells (**Figure 3C and 3D**). In contrast, PAR-binding–deficient mutants exhibited impaired rescue of chromatin condensation, with R93Q showing a near-complete defect and R63A displaying a more modest impairment (**Figure 3E**). These results indicate that efficient PAR-dependent TEAD recruitment is required to stabilize damaged chromatin. Neutral comet assays further showed that wild-type TEAD4 suppressed doxorubicin-induced DNA breaks, whereas PAR-binding–deficient mutants (R63A, R93Q) and the DNA-binding–defective mutant S100A failed to do so (**Figure 3F**).

**Figure 3.**
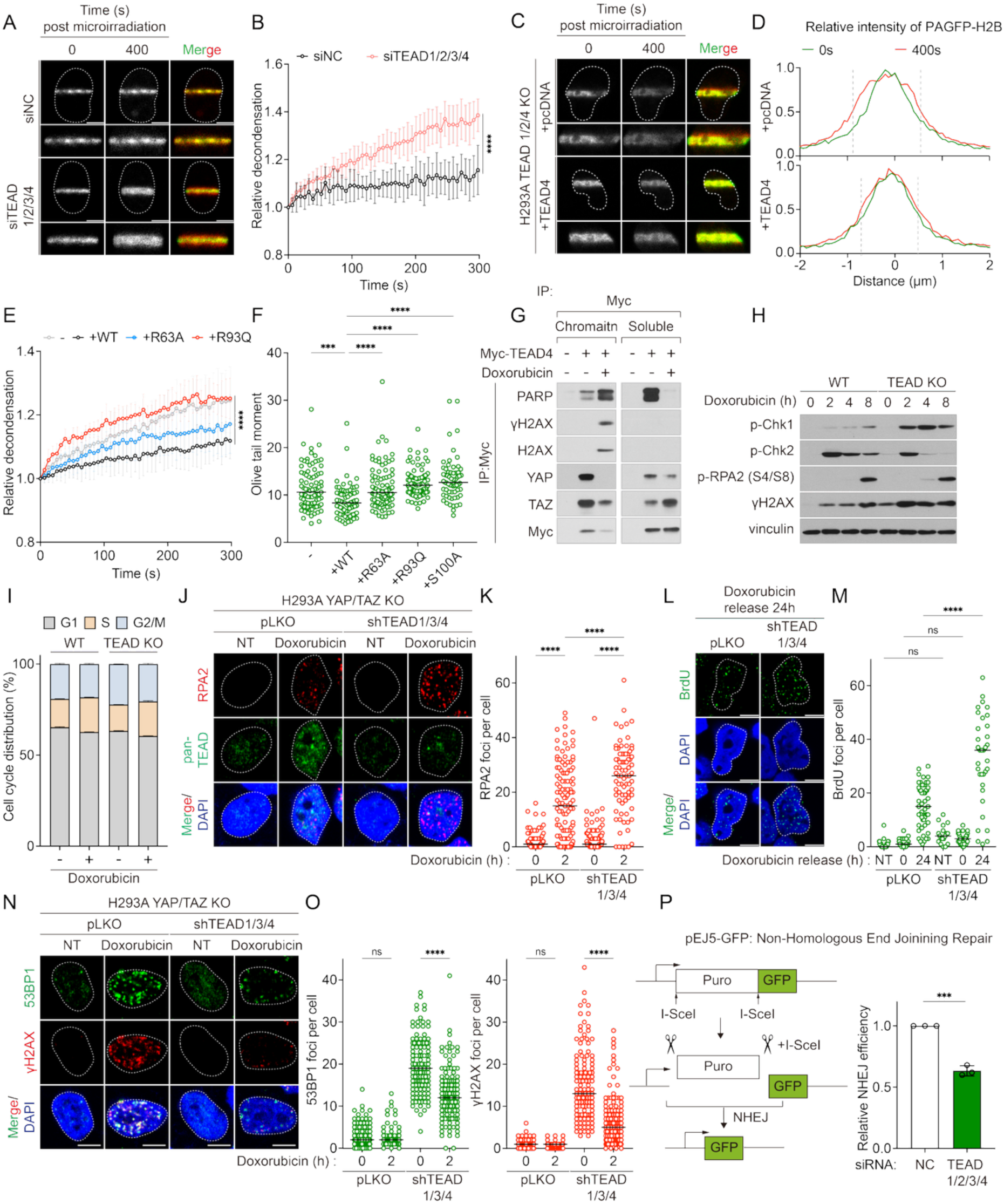
PAR-dependent recruitment of TEAD constrains chromatin relaxation and promotes NHEJ. (A) Confocal images of PAGFP–H2B in U2OS cells transfected with non-targeting control siRNA (siNC, 20 nM) or pooled siTEAD1/2/3/4 (5 nM each), shown at 0 s and 400 s after laser microirradiation. Merged images overlay 0 s (green) and 400 s (red). (B) Quantification of PAGFP–H2B signal width normalized to the 0-s time point (mean ± SD; siNC, n = 7; siTEAD1/2/3/4, n = 10). (C) Confocal images of PAGFP–H2B in TEAD1/2/4-knockout cells reconstituted with mCherry–TEAD4 WT, shown at 0 s and 400 s after microirradiation. Merged images overlay 0 s (green) and 400 s (red). (D) Line-scan analysis of PAGFP–H2B fluorescence intensity at 0 s (green) and 400 s (red) from (C). (E) Quantification of PAGFP–H2B signal width normalized to 0 s in TEAD1/2/4-knockout cells transfected with empty vector (n = 15), mCherry–TEAD4 WT (n = 6), R63A (n = 9), or R93Q (n = 6) (mean ± SD). Data were pooled from two independent biological experiments. (F) Neutral comet assay showing Olive tail moments in HEK293A TEAD1/2/4-knockout cells reconstituted with empty vector, TEAD4 WT, R63A, R93Q, or S100A following doxorubicin treatment (1 μM, 2 h). Sample sizes: empty vector (n = 87), WT (n = 72), R63A (n = 86), R93Q (n = 90), S100A (n = 70). Data were pooled from two independent biological replicates; lines indicate medians. (G) Chromatin immunoprecipitation of Myc–TEAD4 from HEK293A cells treated with doxorubicin showing association with PARP and γH2AX, accompanied by dissociation from YAP/TAZ specifically in chromatin fractions. (H) Immunoblot analysis of DNA damage response signaling in wild-type and TEAD1/2/4-knockout HEK293A cells treated with doxorubicin (5 μM) for the indicated durations. (I) Cell-cycle profiles of HEK293A wild-type and TEAD1/2/4-knockout cells treated with or without doxorubicin (1 μM, 3 h). (J) Confocal images of RPA2 (red) and pan-TEAD (green) in YAP/TAZ-knockout HEK293A cells transduced with control shRNA (pLKO) or shTEAD1/3/4, untreated or treated with doxorubicin (1 μM). (K) Quantification of RPA2 foci per cell from (J). Lines indicate medians (pLKO untreated, n = 133; pLKO + doxorubicin, n = 123; shTEAD untreated, n = 93; shTEAD + doxorubicin, n = 80). (L) Representative confocal images of native BrdU staining (green) in YAP/TAZ-knockout HEK293A cells transduced with pLKO or shTEAD1/3/4, collected 24 h after release from doxorubicin treatment (1 μM, 2 h). (M) Quantification of BrdU foci per cell from (K) (pLKO: NT (n = 49), 0 h release (n = 40), 24 h release (n = 53); shTEAD1/3/4: NT (n = 18), 0 h release (n = 34), 24 h release (n = 35)). Data were pooled from two biological replicates; lines indicate medians. (N) Immunofluorescence images of 53BP1 (green) and γH2AX (red) in YAP/TAZ-knockout HEK293A cells infected with pLKO or shTEAD1/3/4, untreated or treated with doxorubicin (1 μM). (O) Quantification of 53BP1 and γH2AX foci per cell from (N). Sample sizes: pLKO untreated (n = 126), pLKO + doxorubicin (n = 79), shTEAD untreated (n = 126), shTEAD + doxorubicin (n = 129). Lines indicate medians. (P) Schematic of the EJ5-GFP reporter used to assess non-homologous end joining (NHEJ) efficiency. Bar graph shows the percentage of GFP-positive cells following non-targeting control siRNA or siTEAD1/2/3/4 knockdown in pEJ5-GFP–integrated HEK293A cells. Data represent three independent biological replicates. Unless otherwise indicated, scale bars = 5 μm. Error bars represent SD. Lines indicate medians in dot plots. Statistical significance was determined by one-way ANOVA: ns, not significant; *P < 0.05; **P < 0.01; ***P < 0.001; ****P < 0.0001. See also Figure S4.

Chromatin fractionation followed by immunoprecipitation further revealed that DSB induction promotes a chromatin-specific interaction between TEAD and PARP1, coincident with TEAD dissociation from YAP/TAZ (**Figure 3G**). This damage-induced interactome shift positions TEAD within the PARP-dependent chromatin regulatory context at sites of DNA damage. Under these condition, TEAD1/2/4 knockout enhanced checkpoint signaling toward CHK1 activation and increased γH2AX accumulation following doxorubicin treatment, without detectable changes in cell-cycle distribution (**Figure 3H and 3I**).

Given that excessive chromatin relaxation promotes genomic instability^4, 48^, we next examined the consequences of TEAD depletion in YAP/TAZ-knockout HEK293A cells to exclude contributions from canonical YAP/TAZ-TEAD transcriptional activity during the DNA damage response^49–52^. TEAD1/3/4 depletion markedly increased RPA foci (**Figure 3J and 3K**), and native BrdU staining revealed elevated ssDNA exposure after damage release (**Figure 3L and 3M**), consistent with enhanced end resection. These results indicate that PAR-dependent TEAD recruitment restrains chromatin over-relaxation and limits aberrant ssDNA formation at sites of DNA damage.

Because restrained resection is required for efficient non-homologous end joining (NHEJ)^38, 53, 54^, we next assessed whether TEAD loss compromises NHEJ assembly. In the FokI-induced DSB system, TEAD depletion significantly reduced RIF1 recruitment to damage sites (**Figure S3G and S3H**). Similarly, TEAD knockdown attenuated γH2AX accumulation and impaired 53BP1 loading after doxorubicin treatment (**Figure 3N and 3O**). Consistently, EJ5-GFP assays showed a marked reduction in end-joining efficiency^55^ (**Figure 3P**). Together, these results indicate that TEAD maintains a chromatin environment that limits excessive resection and supports efficient NHEJ repair.

### Direct recognition of γH2AX mediates TEAD retention at DNA double-strand breaks

To define the mechanism underlying sustained TEAD retention at DNA lesions, we examined whether TEAD engages γH2AX during the late DNA damage response (DDR). Consistent with microirradiation experiments showing ATM-dependent late retention (**Figure 2F and 2G**), ATM inhibition (KU60019) abolished γH2AX formation and concomitantly eliminated TEAD persistence at FokI-induced DSBs (**Figure 4A and 4B**). By contrast, inhibition of ATR (VE-821), DNA-PK (NU-7441), or PARP (olaparib) had minimal effects on γH2AX accumulation or late TEAD occupancy, indicating that TEAD retention specifically requires ATM-driven γH2AX signaling.

**Figure 4.**
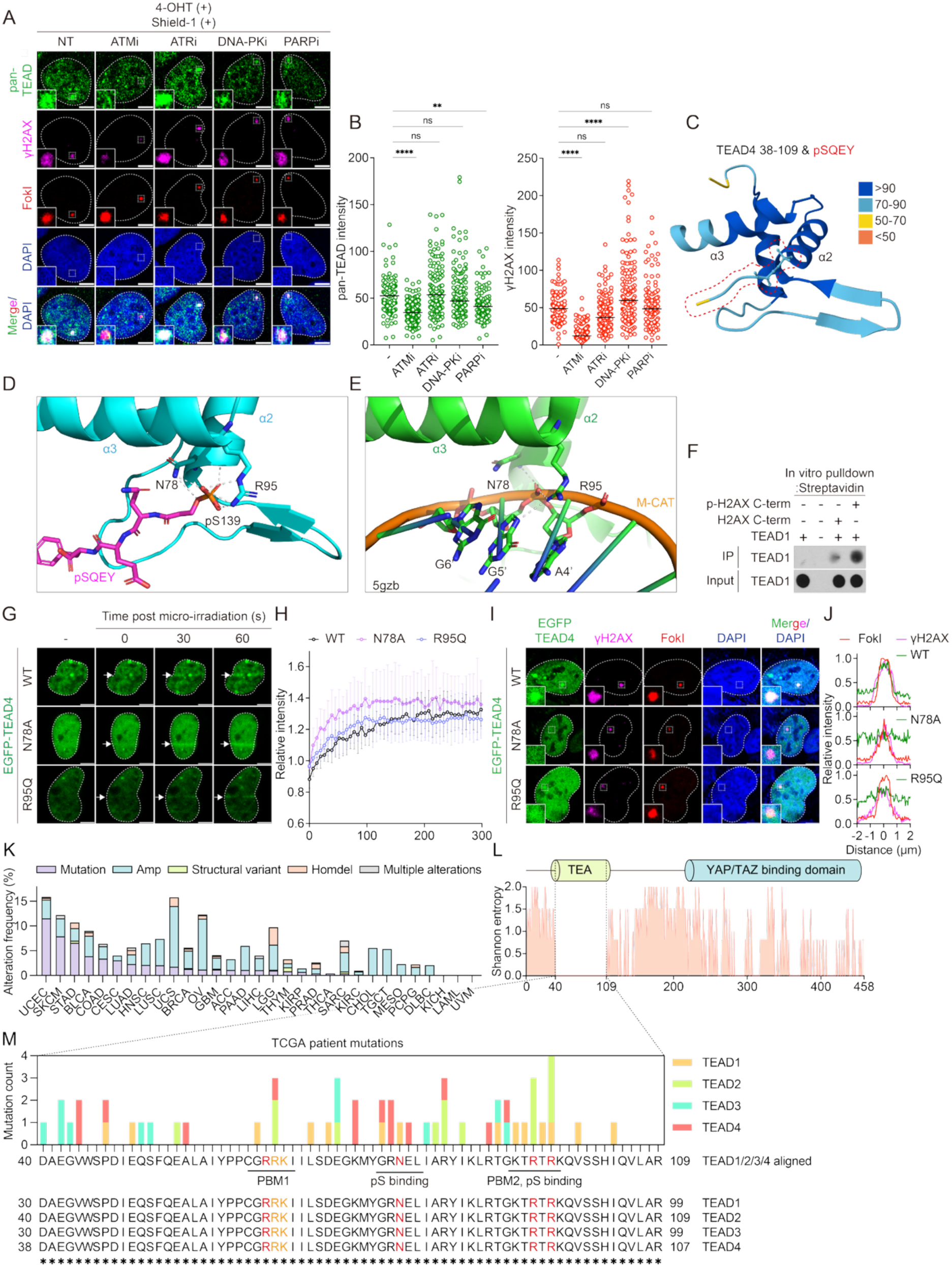
γH2AX-recognition interface in the TEA domain mediates late retention of TEAD at DNA lesions. (A) Confocal images of U2OS 2-6-5 FokI cells treated with vehicle (NT) or the indicated DDR inhibitors—ATM inhibitor (ATMi; KU-60019, 10 μM), ATR inhibitor (ATRi; VE-821, 10 μM), DNA-PK inhibitor (DNA-PKi; NU-7441, 10 μM), or PARP inhibitor (PARPi; olaparib, 10 μM)—for 1 h before induction of a site-specific DSB by 4-OHT and Shield-1. Cells were stained for pan-TEAD (green) and γH2AX (magenta). (B) Dot plots quantifying pan-TEAD and γH2AX fluorescence intensities at FokI-positive versus FokI-negative regions. Lines indicate medians (NT, n = 108; ATMi, n = 165; PARPi, n = 109; ATRi, n = 132; DNA-PKi, n = 136). (C) AlphaFold3 model of TEAD4 (residues 38–109) bound to the phosphorylated H2AX C-terminal tail (pSQEY). Model confidence is indicated by color; α2 and α3 helices of the TEA domain are labeled. (D) Predicted atomic interface showing the phosphate group of γH2AX–Ser139 coordinated by TEAD4 residues N78 (α2 helix) and R95 (α3 helix). Dashed lines indicate predicted hydrogen bonds (TEA domain, cyan; pSQEY peptide, magenta). (E) Comparison of the modeled γH2AX interface (panel D) with the TEAD–DNA crystal structure (PDB: 5GZB), highlighting the analogous phosphate-recognition role of N78 and R95 (TEAD4 in green; M-CAT DNA in orange). (F) In vitro pull-down assay by streptavidin bead showing binding of recombinant TEAD1 to biotinylated H2AX C-terminal peptides (residues 121–143) in either unmodified or phosphorylated (pS139) form. (G) Live-cell imaging of EGFP-TEAD4 (WT, N78A, R95Q) in HEK293A TEAD1/2/4-knockout cells at the indicated times after laser microirradiation. White arrows denote the irradiated region. (H) Quantification of TEAD4 dynamics in (F); fluorescence intensities were normalized to the pre-irradiation signal within the same region (WT, n = 6; R95Q, n = 7; N78A, n = 5). Error bars represent SD. (I) Confocal images of U2OS 2-6-5 FokI cells expressing EGFP-TEAD4 (WT, N78A, R95Q) stained for γH2AX (magenta). (J) Line-scan profiles across the FokI-induced DSB showing relative intensities of EGFP-TEAD4 (green), γH2AX (magenta), and FokI (red), normalized to their respective maximum signals. (K) Alteration frequencies (%) of TEAD1–4 across TCGA cancer types, ordered by mutation rate. Alteration types are indicated as follows: missense/nonsense mutations (purple), amplifications (blue), structural variants (green), homozygous deletions (orange), and multiple alterations (gray). Cancer types are grouped by study and labeled using TCGA abbreviations. (L) Schematic alignment of TEAD1–4 paralogs with corresponding Shannon entropy values, highlighting evolutionarily conserved residues across the TEA domain. (M) Distribution of missense mutations in TEAD1–4 from TCGA pan-cancer datasets. Mutation counts are color-coded by paralog (TEAD1: orange; TEAD2: green; TEAD3: cyan; TEAD4: salmon). Residues experimentally examined in this study are indicated in orange, and those shown to impair recruitment to DNA damage sites are marked in red.

Because the TEA domain lacks canonical ATM phosphorylation motifs (S/TQ)^56^ and we did not detect TEAD phosphorylation after doxorubicin treatment, we hypothesized that TEAD recognizes γH2AX directly. To explore this, we modeled the interaction between the TEA domain of TEAD4 (residues 38-109) and the phosphorylated C-terminal tail of H2AX (pSQEY) using AlphaFold^57^ (**Figure 4C**). The predicted interface positioned the phosphate group on Ser139 of γH2AX in proximity to two conserved TEAD residues, N78 in the α2 helix and R95 in the α3 helix, in a configuration similar to that by which the TEA domain coordinates the DNA phosphate backbone in the published TEAD–DNA crystal structure^16^ (**Figure 4D and 4E**).

In vitro pull-down assays supported this prediction, demonstrating direct binding between TEAD and the phosphorylated H2AX tail (**Figure 4F**). To validate the structural prediction of this interface, we generated TEAD4 N78A and R95Q mutants predicted to disrupt phospho-recognition. Both mutants exhibited normal or slightly accelerated early recruitment, consistent with their intact PAR-binding capacity, but failed to localize to FokI-induced DSBs (**Figure 4G–J**). Together, these findings identify a γH2AX-recognition interface within the TEA domain and establish that TEAD’s late retention at damaged chromatin depends on direct phospho-H2AX binding, mechanistically distinct from its early PAR-dependent recruitment.

### Pan-cancer mutational analysis identifies recurrent alterations in TEAD damage-recognition interfaces

We next examined whether tumor genomes recurrently alter residues within the TEA domain required for TEAD’s DNA damage engagement. Pan-cancer profiling of TEAD1–4 missense mutations revealed recurrent alterations in uterine cancers (UCEC and UCS; >15% overall alterations) and substantial missense burdens in UCEC (11.5%) and SKCM (7.9%) (**Figure 4K**). Structural mapping demonstrated clustering of patient-derived variants within regions corresponding to the PAR-binding interface in the L1 loop and α3 helix (alignment positions 65–67 and 95–97), as well as the γH2AX-recognition surface defined here (alignment positions 80 and 97) (**Figure 4L and 4M**). Notably, recurrent hotspots overlapped residues required for TEAD recruitment to DNA double-strand breaks, indicating that DNA damage–associated interaction surfaces of TEAD are recurrently altered in human cancers.

### Noncanonical TEAD dependency under genotoxic stress in YAP/TAZ-inactive cancers

Having established that TEAD directly engages damaged chromatin through PAR- and γH2AX-dependent mechanisms, we sought to determine whether this activity defines a context-specific functional dependency in tumor cells independent of YAP/TAZ.

Analysis of DepMap datasets identified tumor contexts in which canonical TEAD transcription is inactive, characterized by low YAP/TAZ expression despite sustained TEAD expression^24, 58–60^ (**Figure S4A and S4B**). In these settings, CRISPR dependency profiling revealed minimal reliance on YAP, TAZ, or TEAD paralogs, in contrast to the strong YAP dependency observed in tumors harboring upstream Hippo pathway alterations^61^ (**Figure S4C and S4D**). Consistent with this, TEAD chromatin association and transcriptional output were markedly reduced, and TEAD depletion had negligible effects on basal proliferation (**Figure 5A–D and S4E**), establishing a transcriptionally inactive state.

**Figure 5.**
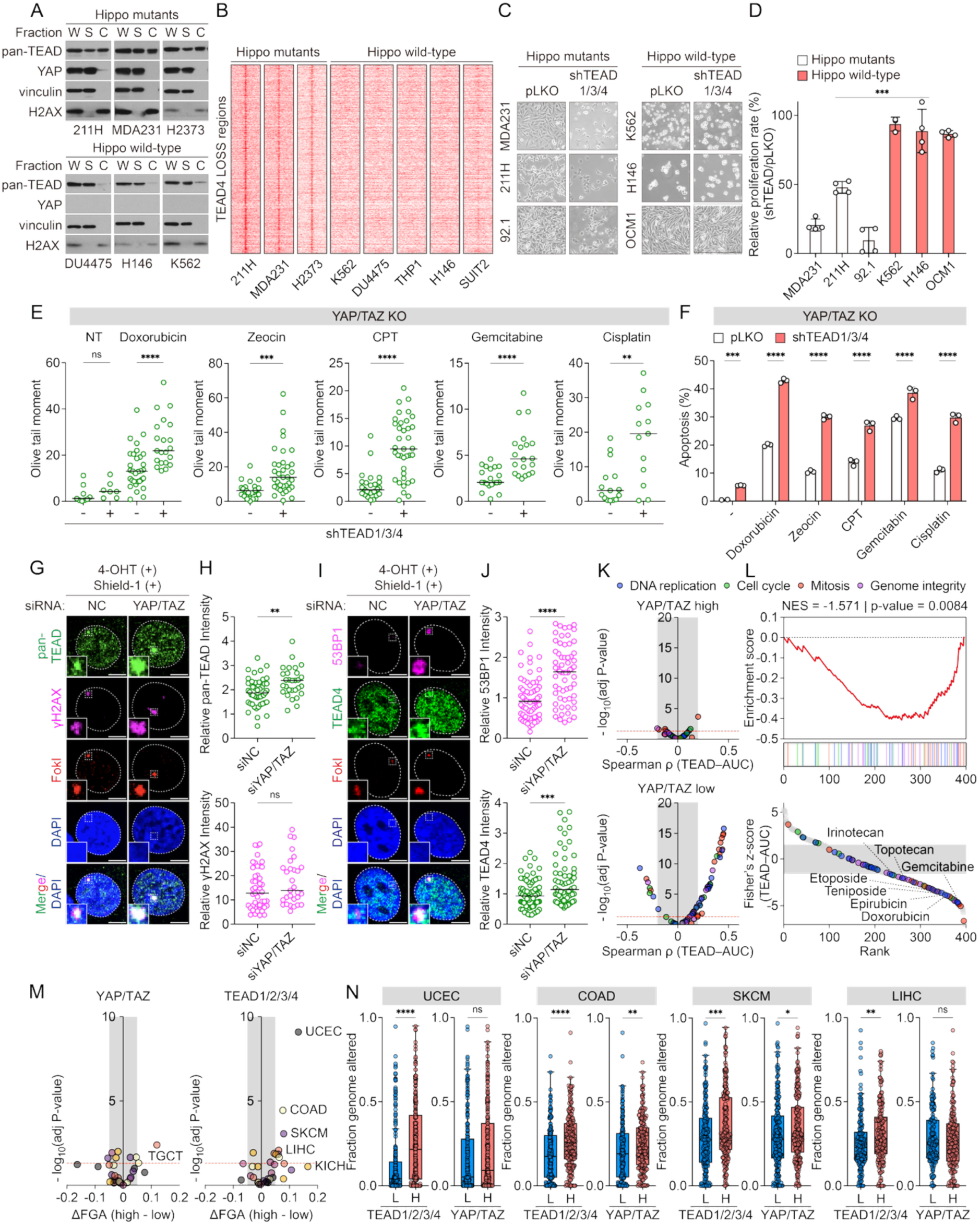
TEAD promotes YAP/TAZ-independent DNA damage tolerance and therapeutic resistance. (A) Immunoblot analysis of chromatin fractionation in Hippo-mutant cancer cell lines (MSTO-211H, MDA-MB-231, H2373) and Hippo–wild-type/YAP/TAZ-low cancers (DU4475, H146, K562). W: whole cell lysate; S: soluble fraction; C: chromatin fraction. (B) TEAD4 ChIP-seq signal at TEAD-LOSS regions (defined in panel C) in Hippo-mutant lines (MSTO-211H, MDA-MB-231, H2373) compared with Hippo–wild-type/YAP/TAZ-low cancers (K562, DU4475, THP-1, H146, SUIT2). (C) Brightfield images of Hippo-mutant cancer cell lines (MDA-MB-231, MSTO-211H, 92.1) and Hippo–wild-type or YAP/TAZ-low lines (K562, H146, OCM-1) transduced with control shRNA (pLKO) or shTEAD1/3/4. (D) Quantification of relative proliferation rates measured by MTT assay 4 days after transduction with pLKO or shTEAD1/2/3/4. Biological replicates: MDA-MB-231, 211H, 92.1, H146, and OCM1 (n = 4 each); K562 (n = 3). (E) Olive tail moments from neutral comet assays in HEK293A YAP/TAZ-knockout cells transduced with pLKO or shTEAD1/2/4 and treated with DNA-damaging agents: doxorubicin (1 μM for 2.5h), zeocin (500 μg/mL for 2.5h), camptothecin (CPT; 2 μM for 6 h), gemcitabine (1 μM for 2.5h) or cisplatin (10 μM for 6 h). Sample sizes: doxorubicin (pLKO: n = 9 untreated, n = 31 treated; shTEAD: n = 7 untreated, n = 21 treated); zeocin/CPT/gemcitabine/cisplatin (pLKO: n = 24, 28, 20, 15; shTEAD: n = 35, 37, 20, 13). Unpaired two-tailed t-tests were used for statistical comparisons. (F) Quantification of apoptosis by PI staining in H293A YAP/TAZ-knockout cells 40 h after release from treatment with the indicated DNA-damaging agents (doxorubicin 1 μM, gemcitabine 1 μM, zeocin 500 μg/mL for 4 h; CPT 2 μM, cisplatin 10 μM for 8 h). Data represent pooled measurements from three independent biological replicates. (G) Recruitment of pan-TEAD and γH2AX to FokI-induced DNA double-strand breaks (DSBs) in U2OS 2-6-5 cells transfected with non-targeting control siRNA (siNC, 20 nM) or combined siYAP (10 nM) and siTAZ (10 nM). (H) Quantification of pan-TEAD and γH2AX fluorescence intensities at FokI-positive regions relative to FokI-negative regions. Lines indicate medians (negative, n = 47; siYAP/TAZ, n = 29). (I) Recruitment of 53BP1 and TEAD4 to FokI-induced DNA double-strand breaks (DSBs) in U2OS 2-6-5 cells transfected with non-targeting control siRNA (siNC, 20 nM) or combined siYAP (10 nM) and siTAZ (10 nM). (J) Quantification of 53BP1 and TEAD4 fluorescence intensities at FokI-positive regions, normalized to siNC. Lines indicate medians (siNC, n = 47; siYAP/TAZ, n = 29). (K) Volcano plots showing the association between mean TEAD1–4 expression and drug response (AUC) across YAP/TAZ-high (top) and YAP/TAZ-low (bottom) cancer cell lines. Each point represents a compound, colored by target pathway: DNA replication (blue), cell cycle (green), mitosis (orange), and genome integrity (purple). The shaded region denotes weak correlations (|ρ| < 0.2), and the red dashed line indicates adjusted P < 0.05. (L) Enrichment of DNA damage–related compounds across drugs ranked by Fisher’s z-transformed differences in TEAD–AUC correlations (YAP/TAZ high − low). The running enrichment score (ES) is shown, with normalized enrichment score (NES) and nominal p-value indicated (Top panel). Ranked distribution of drugs by Fisher’s z, with DNA damage–related compounds highlighted and representative agents labeled. More negative values indicate stronger TEAD-associated drug responses in YAP/TAZ-low contexts (Bottom panel). (M) Volcano plots showing the association between mean YAP/TAZ expression (left) or TEAD1–4 expression (right) and fraction of genome altered (FGA) across cancer types. Each point represents a tumor type. The shaded region denotes small effect sizes (|ΔFGA| < 0.05), and the red dashed line indicates adjusted P < 0.05. Significant cancer types are annotated. (N) Fraction of genome altered (FGA) in TCGA tumors stratified by TEAD1–4 or YAP/TAZ expression (high vs. low, median split). The top nine tumor types showing a significant difference between TEAD-high and TEAD-low groups are shown (UCEC, COAD, SKCM, LIHC). TEAD or YAP/TAZ-low group n = 255/210/216/177; TEAD or YAP/TAZ-high group n = 255/211/217/178.

Despite this, TEAD became functionally essential under genotoxic stress. TEAD depletion in YAP/TAZ-deficient cells led to increased DNA damage accumulation and apoptosis following treatment with double-strand break–inducing agents (**Figure 5E-F and S4F**), indicating that TEAD supports DNA damage tolerance independently of its canonical transcriptional role. Mechanistically, YAP/TAZ suppression enhanced TEAD recruitment to DNA damage sites and promoted accumulation of the NHEJ factor 53BP1 without altering γH2AX levels, consistent with activation of a transcription-independent DNA repair axis (**Figure 5G–J**).

### TEAD-dependent DNA damage tolerance drives therapeutic resistance and genomic instability

This context-dependent requirement for TEAD was reflected in therapeutic response. In GDSC datasets^62^, TEAD expression correlated with resistance to DNA-damaging agents specifically in YAP/TAZ-low cancers, with the strongest associations observed for double-strand break–inducing compounds (**Figure 5K and 5L**). These findings support a role for TEAD-dependent DNA damage tolerance in promoting survival under genotoxic stress in transcriptionally inactive contexts.

Extending this analysis to human tumors, TEAD expression was selectively associated with genomic instability across 10,967 TCGA samples, whereas YAP/TAZ expression, Hippo target gene signatures failed to effectively stratify tumors with elevated genomic instability (**Figure 5M and S4G**). In multiple cancer types, including UCEC, COAD, SKCM, LIHC, BLCA, LUAD, and HNSC, TEAD-high tumors exhibited significantly increased FGA and poorer clinical outcome, particularly among patients receiving genotoxic therapies (**Figure 5N and S4H,I**). Together, these findings suggest that TEAD-dependent DNA damage tolerance contributes to therapeutic resistance and may support the fitness of genomically unstable tumors, linking a transcription-independent DNA repair program to cancer progression.

### Allosteric inhibition of the TEAD N-terminus disrupts its noncanonical DNA repair function

Given the mechanistic and clinical evidence that TEAD DNA damage function is essential and YAP/TAZ-independent, we sought to selectively inhibit its N-terminal DDR interface. We screened an in-house chemical library for compounds capable of binding TEAD^63^. BY116 was identified through screening for TEAD-binding compounds that preserved YAP/TAZ association. Immunoprecipitation of TEAD showed that BY116 perturbed N-terminal TEAD associations, including TEAD–PARP interaction, without disrupting YAP/TAZ binding (**Figure 6A**). Consistent with targeting the N-terminal domain, this compound series reduced TEAD-dependent NHEJ activity in vitro (**Figure 6B**).

**Figure 6.**
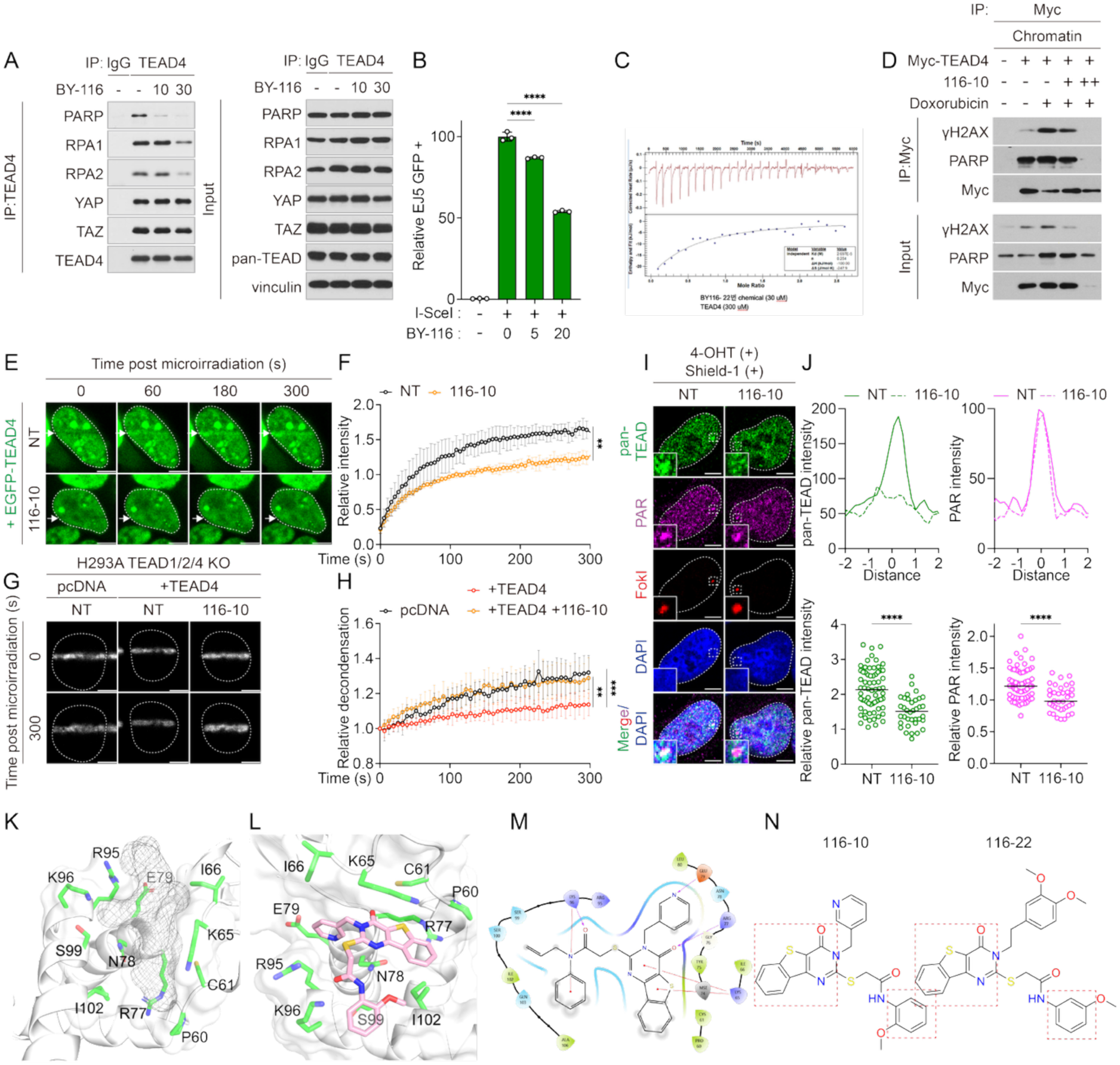
Allosteric targeting of the TEAD N-terminus disrupts its noncanonical DNA repair function. (A) Immunoblot analysis of endogenous TEAD4 immunoprecipitated in HEK293A cells treated with BY116 (10 or 30 μM, 18 h). (B) EJ5-GFP reporter assay measuring NHEJ efficiency in cells transfected with I-SceI for 48 h and treated with 116-10 (5 or 20 μM, 24 h). Data represent three independent biological replicates. (C) Isothermal titration calorimetry (ITC) measuring binding affinity (KD) of BY116-22 derivatives to TEAD. (D) Immunoblot of Myc–TEAD4 immunoprecipitated from chromatin fractions of HEK293A TEAD1/2/4-knockout cells expressing Myc–TEAD4 and treated with doxorubicin (5 μM, 2 h) in the presence or absence of 116-10 (10 μM (+) or 30 μM (++), 24 h). (E) Live-cell imaging of EGFP–TEAD4 in HEK293A TEAD1/2/4-knockout cells pretreated with 116-10 (10 μM, 18 h) prior to microirradiation. White arrows indicate irradiated regions. (F) Quantification of EGFP–TEAD4 recruitment dynamics at microirradiated tracks, normalized to pre-irradiation intensity (mean ± SD; NT, n = 3; 116-10, n = 3). (G) Confocal images of PAGFP–H2B at 0 s and 300 s in HEK293A TEAD1/2/4-knockout cells transfected with empty vector or mCherry–TEAD4 and treated with 116-10 (20 μM, 8 h). (H) Quantification of PAGFP signal width normalized to the 0-s time point (mean ± SD; empty vector, n = 14; TEAD4, n = 12; TEAD4 + 116-10, n = 10). (I) Confocal images showing recruitment of pan-TEAD and PAR to FokI-induced DSBs in U2OS 2-6-5 cells treated with 116-10 (20 μM, 8 h). (J) Line-scan profiles of pan-TEAD (green) and PAR (magenta) signals with or without 116-10 at FokI-positive regions (n = 3 per condition, top), and quantification of fluorescence intensities at FokI-positive regions normalized to FokI-negative regions (NT, n = 68; 116-10, n = 37; medians shown). (K) Predicted binding pocket 1 identified by PASSer2.0. Residues within 4 Å of pocket 1 are shown. Amino acids are displayed as surface representations colored using a rainbow scheme. (L) Predicted binding mode of compound 116-10 (pink) within the TEAD4 N-terminal allosteric pocket. (M) Interaction diagram of compound BY116 with TEAD4. Red lines indicate π–cation interactions, and purple arrows indicate hydrogen bonds. (N) Chemical structures of 116-10 and 116-22. The common structural ring scaffold is highlighted with a red dashed box. Unless otherwise indicated, scale bars = 5 μm. Statistical significance was calculated by one-way ANOVA: ns, not significant; *P < 0.05; **P < 0.01; ***P < 0.001; ****P < 0.0001.

To enable direct biophysical validation and improve cellular applicability, we generated a set of derivatives retaining the core scaffold. Among these, BY116-22 was suitable for quantitative binding analysis and exhibited measurable interaction with the TEAD N-terminus by isothermal titration calorimetry (ITC), confirming direct target engagement (**Figure 6C**). In parallel, BY116-10 was optimized for cellular assays and was therefore used to evaluate inhibition in the DNA damage context. BY116-10 effectively suppressed damage-induced TEAD chromatin engagement, demonstrating functional inhibition of the DDR-associated TEA domain interface (**Figure 6D**). Functionally, BY116-10 phenocopied genetic disruption of TEAD’s DDR interface. Treatment suppressed early TEAD recruitment to laser-induced double-strand breaks (**Figure 6E and 6F**), impaired TEAD-dependent chromatin condensation (**Figure 6G and 6H**), and markedly reduced TEAD accumulation at FokI-induced breaks, despite only a modest reduction in PAR formation (**Figure 6I and 6J**), consistent with inhibition of both early recruitment and late retention mechanisms.

Structural modeling provided a rationale for these inhibitory effects. PASSer2.0 predicted allosteric N-terminal pockets (**Figure 6K and S5A**) and docking analyses positioned BY116 and BY116-10 within this cavity adjacent to residues required for PAR and γH2AX recognition^64, 65^ (**Figure 6L and S5B**). The conserved aromatic scaffold formed stabilizing interactions within this pocket, a feature retained in BY116-10 and BY116-22 (**Figure 6M and 6N**). AlphaFold3 modeling independently supported the predicted N-terminal pocket as a druggable site (**Figure S5C**), and molecular dynamics simulations were consistent with stable engagement of this interface (**Figure S5D-F**).

Together, these findings establish that the TEAD N-terminal DDR interface is pharmacologically targetable. Allosteric inhibition disrupts TEAD’s DNA damage program and phenocopies genetic TEAD disruption, supporting a therapeutic strategy for Hippo–wild-type and YAP/TAZ-independent cancers.

### TEAD inhibition sensitizes YAP/TAZ-independent tumors to DSB-inducing chemotherapy

We next examined the therapeutic consequences of inhibiting TEAD’s DDR function in YAP/TAZ-independent tumor models. In YAP/TAZ-low myeloid cancer cells with differential TEAD expression^66, 67^, TEAD abundance predicted sensitivity to combined TEAD inhibition and genotoxic stress. TEAD-low AR230 cells were intrinsically sensitive to doxorubicin and showed minimal additional response to BMY116-10, whereas TEAD-high K562 cells exhibited significant drug synergy between doxorubicin and BMY116 as quantified by Bliss independence analysis (**Figure 7A and S6A**). Re-expression of TEAD in AR230 restored damage-induced recruitment of TEAD and associated repair factors to chromatin, which was suppressed by BMY116-10 (**Figure 7B**).

**Figure 7.**
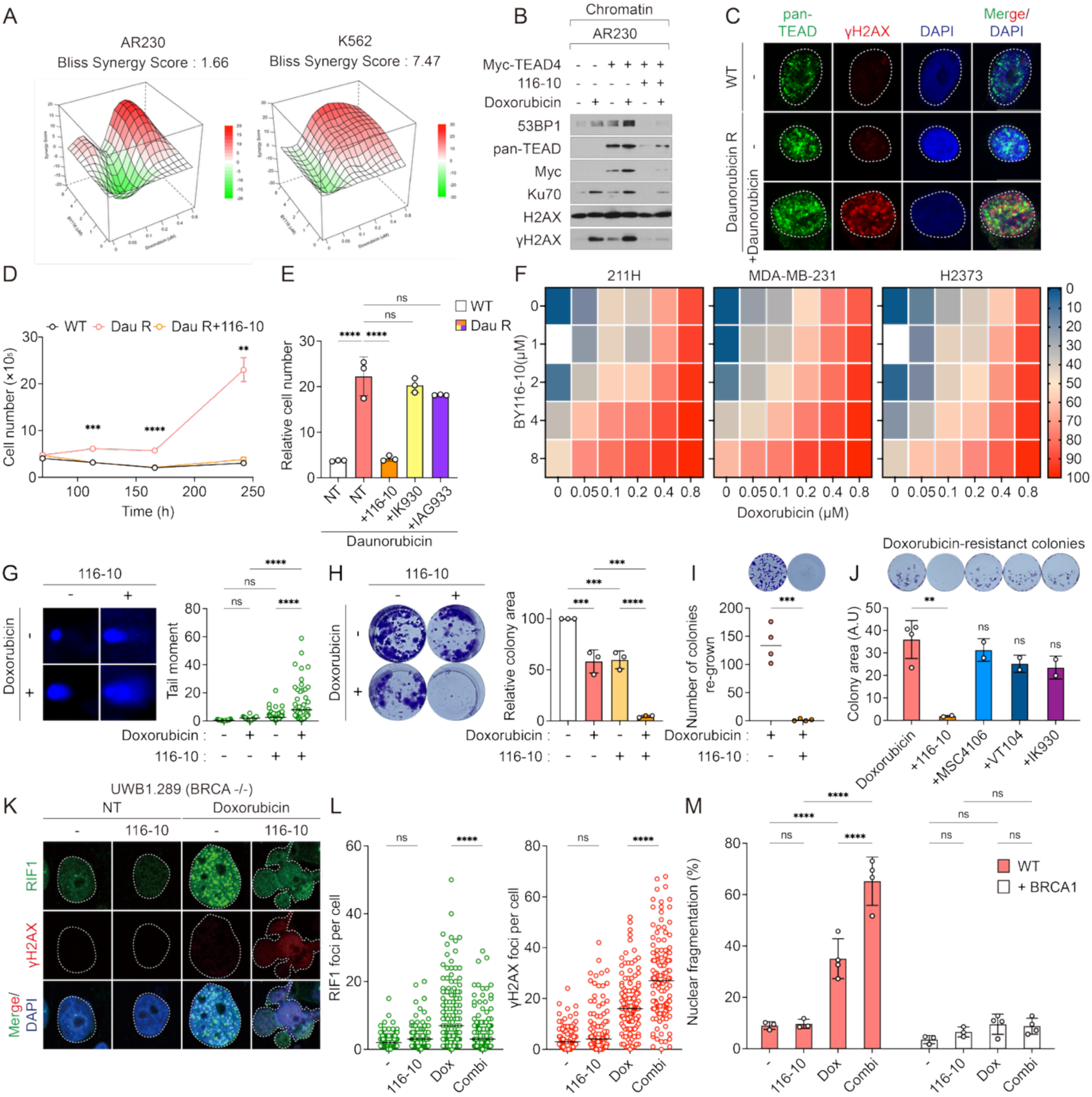
TEAD inhibition exploits DNA repair vulnerabilities to sensitize tumors to chemotherapy. (A) Bliss surface plot showing drug synergy between 116-10 and doxorubicin in TEAD-high K562 cells (synergy score = 7.47) but not in AR230 cells (synergy score = 1.66). (B) Immunoblot analysis of chromatin fractions from AR230 cells expressing Myc-TEAD4 treated with doxorubicin (1 μM, 6 h) with or without 116-10 (20 μM, 18 h). (C) Immunofluorescence images of U937 wild-type (NT) and daunorubicin-resistant (DauR) cells (NT or daunorubicin-treated) stained with pan-TEAD (green) and γH2AX (red). (D) Survival curve of U937 cells treated with daunorubicin (10 nM) for 244 h. Cell number was calculated from ATP luminescence measured by CellTiter-Glo. (E) Relative cell number at 240 h in U937 wild-type and daunorubicin-resistant cells treated with NT, 116-10, IK930, or IAG933 (1 μM). (F) Bliss synergy maps for 116-10 (1, 2, 4, 8 μM) in combination with doxorubicin (0.05, 0.1, 0.2, 0.4, 0.8 μM) for 72 h in MSTO-211H, MDA-MB-231, and H2373 cells. (G) Neutral comet assay in H2373 cells treated with doxorubicin (1 μM) with or without 116-10 (10 μM). (H) Clonogenic survival assay in H2373 cells following treatment with doxorubicin (1 μM) with or without 116-10 (10 μM). (I) Colony growth of doxorubicin-resistant clones in the presence or absence of 116-10. (J) Relative colony area of doxorubicin-resistant clones treated with 116-10, MSC4106, VT104, or IK930 (all 1 μM), compared to untreated controls. (K) Immunofluorescence images of UWB1.289 (BRCA-/-) cells treated with doxorubicin and 116-10, stained for RIF1 (green) and γH2AX (red). (L) Quantification of foci per cell in (K). Black lines indicate medians. (M) Quantification of nuclear fragmentation in cells treated with doxorubicin and/or 116-10.

Given that tolerance to DSB-inducing chemotherapy promotes survival under sustained treatment, we examined whether TEAD contributes to acquired chemoresistance in acute myeloid leukemia (AML), a malignancy primarily treated with DNA-damaging agents^68,69^. In U937 cells, prolonged daunorubicin exposure led to the emergence of resistant clones accompanied by marked upregulation of endogenous TEAD (**Figure S6B and S6C**). Elevated TEAD co-localized with γH2AX following daunorubicin treatment, consistent with engagement of the DNA damage program (**Figure 7C**).

Pharmacological inhibition with BMY116 abrogated resistance, whereas a canonical YAP/TAZ–TEAD inhibitor (IK930, IAG933) had no effect^70, 71^, indicating that chemoresistance depends specifically on TEAD’s DDR function rather than transcriptional activity (**Figure 7D and 7E**). Consistent with this model, analysis of chemotherapy-treated AML patient cohorts showed that high TEAD expression correlated with poor clinical outcome, whereas YAP/TAZ levels lacked prognostic value (**Figure S6D**). Together, these findings support a role for TEAD in acquired chemoresistance and demonstrate that selective DDR inhibition can re-sensitize resistant AML cells to genotoxic therapy.

Finally, we extended TEAD DDR inhibition to Hippo-mutant and Hippo–wild-type cancers^60^. Although Hippo-mutant tumors are canonically YAP/TAZ-dependent, DSB induction triggered rapid YAP/TAZ dissociation and transient suppression of canonical target genes, indicating functional disengagement from transcription (**Figure 1J and S6E**). Under these conditions, combined treatment with doxorubicin and BY116-10 produced significant synergy across multiple Hippo-mutant models, whereas a classical YAP/TAZ–TEAD inhibitor showed only modest effects (**Figure 7F and S6F**). Comet assays revealed increased DNA break accumulation and apoptosis following combination treatment, consistent with impaired DSB repair (**Figure 7G and S6G and S6H**). Long-term clonogenic assays further demonstrated that BY116-10 markedly suppressed colony regrowth after doxorubicin exposure, limiting resistant outgrowth (**Figure 7H and 7I and S6I**).

Importantly, canonical YAP/TAZ inhibitors^72, 73^ failed to suppress doxorubicin-resistant colony outgrowth, indicating that TEAD’s N-terminal DDR activity, rather than its transcriptional function, is required for survival and regrowth under genotoxic stress (**Figure 7J**). In Hippo-wild-type SUIT2 cells, BY116-10 but not canonical YAP/TAZ inhibitors enhanced DNA damage accumulation in response to doxorubicin (**Figure S6J**). Together, these findings demonstrate that pharmacological targeting of TEAD’s DNA damage function enhances chemotherapy efficacy by limiting repair-mediated survival and resistance, thereby proposing a non-transcriptional vulnerability that remains inaccessible to conventional YAP/TAZ-directed inhibitors.

### TEAD inhibition exploits DNA repair vulnerabilities in BRCA-mutant cancers

Because TEAD inhibition compromises NHEJ repair, we asked whether HR-deficient tumors exhibit increased sensitivity to this perturbation. BRCA1-mutant ovarian cancer cells (UWB1.289) and their BRCA1-rescued counterparts provided a genetically defined system to evaluate this vulnerability^74, 75^. In BRCA1-mutant cells, co-treatment with doxorubicin and BY116-10 markedly reduced RIF1 foci and increased γH2AX accumulation and nuclear fragmentation, indicating the persistence of unrepaired DNA damage (**Fig. 7K and 7L**). While BRCA1-rescued cells displayed increased RAD51 foci indicative of restored HR capacity, BRCA1-mutant cells lacked compensatory HR activation and underwent synergistic apoptosis (**Figure 7M and S6K**). These results indicate that disruption of NHEJ by TEAD inhibition is incompatible with HR deficiency. Consequently, TEAD inhibition exposes a repair vulnerability in BRCA-mutant cancers, sensitizing them to DNA double-strand break–inducing agents.

## Discussion

Although transcription factors (TFs) have been implicated in the DNA damage response, their contributions have largely been interpreted through transcriptional activity^52, 76–78^. Here, we identify the oncogenic transcription factor TEAD as a direct regulator of damaged chromatin. Rather than acting solely through gene regulation, TEAD promotes DNA repair, revealing a broader role for transcription factors in genome maintenance. These findings indicate that transcription factors can be repurposed under genotoxic stress as chromatin regulators with functions beyond canonical transcription and relevance for therapeutic targeting.

As chromatin architecture is increasingly recognized as a key determinant of DNA repair efficiency^2, 47, 79^, this gap becomes more apparent. Transcription factors, despite their ability to shape chromatin through sequence-specific DNA binding and diverse protein interactions^31, 80, 81^, have been largely excluded from frameworks of active chromatin regulation during DNA repair^12, 82^. This likely reflects the absence of clear mechanistic frameworks and defined domains linking transcription factors to repair processes. Even in well-studied cases such as p53^13, 14^, transcription-independent roles in DNA damage repair have not been integrated into a broader model of chromatin-based repair.

We show that TEAD mediates DNA repair through a transcription-independent mechanism driven by its TEA domain, separable from its canonical coactivator-dependent activity and directly targetable. Structure-guided prediction combined with experimental validation identifies small molecules that engage TEAD and suppress its DNA repair function, establishing a framework for targeting transcription factors as regulators of chromatin dynamics in cancer.

At the mechanistic level, the TEA domain functions as a DNA damage–responsive chromatin engagement interface. Residues including R63, R93, and R95 mediate biphasic recruitment through PAR-dependent accumulation followed by γH2AX-dependent retention at sites of DNA double-strand breaks. Temporal recruitment has been described for select repair factors^39, 40, 83^; however, coordinated biphasic engagement within a transcription factor has not been clearly defined. TEAD recruitment requires an intact DNA-binding interface, and residues involved in PAR recognition overlap with the canonical DNA-binding surface, indicating that a conserved structural interface is repurposed for damage sensing^11, 16, 44^. Structural modeling further supports this mechanism, suggesting that γH2AX recognition may exploit a binding mode analogous to DNA backbone interactions^57^. Several residues within this recruitment interface overlap with recurrent mutations observed in human tumors. Although the number of cases is limited, this convergence suggests that disruption of TEAD-mediated chromatin engagement may represent an alternative mode of genome maintenance adaptation in cancer.

This mode of chromatin engagement is coupled to rapid chromatin remodeling. TEAD recruitment correlates with chromatin condensation at damage sites, supporting an active role in shaping local chromatin architecture during the DNA damage response^4, 9, 84^. The underlying mechanism remains unresolved, whether through direct interaction with PARylated chromatin or recruitment of additional factors; however, the data support a model in which TEAD functions as a chromatin remodeler rather than a passive DNA-binding factor. TEAD also associates with multiple DNA repair proteins, including PARP1, Ku70/80,

RPA, and Rad51^46, 85, 86^. These interactions are separable from the recruitment interface and do not account for TEAD localization to DNA lesions, indicating that chromatin engagement is not driven solely by protein–protein interactions. DNA damage induces a chromatin-specific interactome shift in which TEAD associates with PARP1 and γH2AX while dissociating from YAP/TAZ, consistent with a functional transition from transcriptional coactivator to a DNA damage–responsive chromatin regulator. The upstream signals governing this switch remain incompletely defined, although partial rescue by ATM inhibition implicates damage-induced signaling in TEAD–YAP dissociation. Whether additional post-translational modifications coordinate this transition remains to be determined.

Loss of TEAD disrupts chromatin compaction at sites of DNA damage, leading to excessive relaxation, ssDNA exposure, and increased RPA loading. This aberrant chromatin state constrains non-homologous end joining (NHEJ) and impairs repair of double-strand breaks, promoting genomic instability. Defects in early TEAD recruitment are sufficient to induce ssDNA accumulation and compromise NHEJ, indicating that TEAD-dependent chromatin organization is required to maintain repair pathway fidelity. TEAD-mediated NHEJ contributes to DNA damage tolerance and chemoresistance, supported by experimental assays and pharmacogenomic analyses linking TEAD function to sensitivity to double-strand break–inducing agents such as doxorubicin, etoposide, and teniposide. TEAD dependency is more pronounced in response to blunt-ended DSBs, suggesting preferential activity in specific structural contexts of DNA damage.

This function is largely independent of canonical Hippo pathway activity. While YAP/TAZ-driven transcription is a major driver of tumor progression^19^, TEAD-mediated DNA repair operates through a distinct mechanism. Patient datasets show that TEAD expression correlates with genomic instability and poor clinical outcome in contexts with low Hippo pathway target gene activity. This relationship is particularly evident in breast cancer, where elevated TEAD expression associates with increased fraction of genome alteration despite reduced YAP/TAZ transcriptional output, indicating that TEAD supports genome maintenance and therapeutic resistance in Hippo-inactive tumors. Targeting this noncanonical function expands the therapeutic landscape of TEAD beyond YAP/TAZ-dependent contexts. Structure-guided identification of a druggable pocket within the TEA domain^65^, combined with experimental validation, enabled the discovery of small molecules that directly engage TEAD and suppress its DNA damage response activity. This strategy provides a framework for targeting transcription factors through noncanonical structural features.

This dependency extends to tumor types not captured by YAP/TAZ-driven models. In acute myeloid leukemia (AML) and small cell lung cancer (SCLC), where YAP/TAZ activity is low yet TEAD expression is maintained^24^, TEAD contributes to chemotherapy resistance. TEAD expression is elevated in daunorubicin-resistant AML cells, and inhibition of the TEA domain restores drug sensitivity, whereas canonical YAP/TAZ–TEAD inhibitors are ineffective. Patient data further support this association, linking TEAD expression to poor outcome following anthracycline-based chemotherapy.

These results indicate that TEAD supports survival under genotoxic stress through a noncanonical DNA damage response function, particularly in Hippo-inactive settings, and suggest that effective therapeutic strategies may require combined or sequential inhibition of both TEAD DNA repair activity and its canonical transcriptional function^67^. Collectively, TEAD undergoes a context-dependent functional transition at sites of DNA double-strand breaks, operating independently of its YAP/TAZ-mediated transcriptional role to regulate chromatin dynamics and DNA repair. More broadly, transcription factors can be repurposed as chromatin regulators during DNA damage, revealing therapeutically targetable functions beyond canonical transcription.

## Acknowledgement

We thank Kun-Liang Guan and Tracy T. Tang for insightful discussions and critical feedback on TEAD inhibition and emerging therapeutic strategies. We also thank Ho-Jae Lee (Leica Microsystems) for assistance with microscopy setup and experimental optimization. The results shown here are in part based upon data generated by the TCGA Research Network (https://www.cancer.gov/tcga).

## Author Contribution

Conceptualization, D.-H.K., H.W.P.; methodology, D.-H.K., J.K., J.-S.R., and D.S.; investigation, D.-H.K., S.K., H.-R.K., J.K., and J.-S.M.; formal analysis, D.-H.K., J.K., S.K., H.- R.K., H.J., E.A.L., H.R.S., H.C.K., K.M., and Y.K.; data curation, J.K., H.-R.K., H.R.S., H.C.K., S.C.C., E.J., J.B.K., K.M., and Y.K.; resources, I.K., Y.K., W.K., and K.T.N.; writing – original draft, D.-H.K. and J.K.; writing – review & editing, D.-H.K., J.K., H.J., H.-R.K., J.-S.R., D.S., K.T.N., and H.W.P.; supervision, J.-S.R., D.S., K.T.N., and H.W.P.; funding acquisition, H.W.P.

## Method

### Cell lines

HEK293A, HEK293A YAP/TAZ knockout (KO), LATS1/2 KO, TEAD1/2/4 KO, U2OS, U2OS 2-6-5, and MDA-MB-231 cells were cultured in DMEM (Hyclone, SH30022.01) supplemented with 10% fetal bovine serum (FBS; Hyclone, SV30207.02) and 1% penicillin–streptomycin (P/S; Invitrogen, 15140122). The HEK293A CRISPR KO cell lines were kindly provided by Dr. Kun-Liang Guan, and U2OS 2-6-5 reporter cells were generously provided by Dr. Roger A. Greenberg. SUIT2, 92.1, H2373, MSTO-211H, K562, AR230, Molm13, Molm14, OCM1, and H146 cells were maintained in RPMI-1640 (Hyclone, SH30027.01) containing 10% FBS and 1% P/S. UWB1.289 parental and BRCA1-reconstituted derivatives were cultured in a 1:1 mixture of RPMI-1640 and Mammary Epithelial Cell Growth Medium (MEGM; Lonza, CC-3150) supplemented with 3% FBS and 1% P/S; BRCA1-reconstituted cells were additionally maintained in 200 μg/ml G418 (Geneticin; Gibco, 10131-035). All cell lines were maintained at 37 °C in a humidified incubator with 5% CO₂.

### Transfection

HEK293A, HEK293A YAP/TAZ KO, HEK293A TEAD1/2/4 KO, HEK293A LATS1/2 KO, U2OS, and U2OS 2-6-5 cells were transfected with plasmid DNA using jetPRIME® reagent (Polyplus). Plasmid DNA was incubated with jetPRIME® for 30 min to allow complex formation and then added to cells at 60–70% confluency. Cells were used for experiments 48– 72 h post-transfection. For siRNA transfection, Lipofectamine RNAiMAX (Invitrogen, 13778075) was used according to the manufacturer’s instructions. siRNAs were transfected at a final concentration of 20 nM. A non-targeting Dharmacon control siRNA (#2, D-001210-02-20) was used as the negative control. The following siRNAs were used: siYAP1 (M-012200-00-0020), siWWTR1 (M-016083-00-0020), siTEAD1 (M-012603-01-0020), siTEAD2 (M-012611-00- 0020), siTEAD3 (M-012604-01-0020), and siTEAD4 (M-019570-03-0020).

### Immunoprecipitation

Cells were lysed in ice-cold 0.5% NP-40 lysis buffer (25 mM Tris-HCl, pH 7.6; 150 mM NaCl; 0.5% NP-40; 50 U/mL Turbonuclease (Sigma-Aldrich, T4330)) supplemented with protease inhibitors. Lysates were pipetted 20–30 times every 10 min for 30 min on ice and centrifuged at 13,000 rpm for 10 min at 4 °C. For endogenous protein immunoprecipitation, clarified supernatants were incubated overnight at 4 °C with primary antibody (TEAD4; TEF-3 Antibody (N-G2), sc-101184; 400 ng), followed by incubation with Protein G magnetic beads (Pierce™ Protein G Magnetic Beads, 88848) for 2 h at 4 °C. For Myc-tagged proteins, supernatants were incubated overnight at 4 °C with anti-c-Myc magnetic beads (Pierce™ Anti-c-Myc Magnetic Beads, 88843). Immunocomplexes were washed four times with NP-40 lysis buffer and eluted in 2× Laemmli buffer for SDS–PAGE.

For chromatin-fraction immunoprecipitation, soluble (supernatant) and chromatin-enriched (pellet) fractions were first isolated by chromatin fractionation. Each fraction was incubated overnight at 4 °C with anti-c-Myc magnetic beads (Pierce™ 88843), washed four times with NETN buffer, and eluted in 2× Laemmli buffer for SDS–PAGE.

### In vitro pulldown assay

To evaluate TEAD–poly(ADP-ribose) (PAR) interaction, recombinant PAR (4 ng; Bio-Techne #4336-100-01) and TEAD1 (4 ng; LSbio #LS-G57536) were incubated for 30 min at room temperature in incubation buffer (25 mM Tris-HCl, pH 7.6, 150 mM NaCl) supplemented with protease inhibitors. TEAD1 was subsequently immunoprecipitated, washed with buffer supplemented with 0.5% NP-40, and associated PAR was detected by immunoblotting using an anti-PAR antibody.

To assess TEAD–γH2AX interaction, biotinylated peptides corresponding to the H2AX C-terminal region (amino acids 130–142; TSATVGPKAPSGGKKATQASQEY) were synthesized in both unmodified and phosphorylated forms. Recombinant TEAD1 (4 ng) was incubated with H2AX or phospho-H2AX peptides (8 ng) for 30 min at room temperature under the same buffer conditions. Complexes were captured using streptavidin magnetic beads, washed with buffer containing 0.5% NP-40, and analyzed by immunoblotting for TEAD1. Wash conditions and handling steps were consistent with those used for immunoprecipitation assays.

### Chromatin fractionation

Cells were lysed in ice-cold NETN buffer (20 mM Tris-HCl, pH 7.6, 100 mM NaCl, 1 mM EDTA, 0.5% NP-40) supplemented with Halt™ Protease and Phosphatase Inhibitor Cocktail (Thermo). Lysates were incubated on ice for 25 min and centrifuged at 13,000 rpm for 10 min at 4 °C. The supernatant was collected as the soluble fraction. The pellet was washed twice with NETN buffer, resuspended in NETN containing TurboNuclease (50 U/mL), and incubated for 30 min at room temperature to release chromatin-associated proteins. Both soluble and chromatin fractions were denatured in 2× Laemmli buffer for SDS–PAGE or processed for immunoprecipitation as described.

For cytosol / nucleoplasm / chromatin fractionation, cytosolic extracts were first obtained using the NE-PER™ Nuclear and Cytoplasmic Extraction Kit (Thermo, N78833) according to the manufacturer’s protocol. After removal of the cytosolic fraction, the remaining insoluble material—containing intact nuclei—was subjected to the subsequent fractionation steps. Nuclear pellets were incubated in NETN buffer on ice for 25 min, and the resulting supernatant was collected as the nucleoplasmic fraction. The remaining pellet was washed and resuspended in NETN supplemented with TurboNuclease (50 U/mL), following the same conditions used for chromatin isolation above, to extract chromatin-associated proteins. All fractions were denatured in 2× Laemmli buffer for SDS–PAGE or used for immunoprecipitation as indicated.

### Chromatin immunoprecipitation (ChIP) assay

For ChIP experiments, cells were trypsinized into single cells to yield 3 x 107 cells. The cells were cross-linked with 1 % formaldehyde for 15 min, quenched with 0.125 M glycine for 10 min at room temperature, and washed with PBS. Cell pellets were lysed in 1,200 μl of cell lysis buffer (10 mM Tris-Cl pH 8.0, 10 mM NaCl, 0.2 % NP-40), supplemented with protease inhibitor (cOmplete™ Protease Inhibitor Cocktail, Cat#11697498001; Roche) and 1 mM DTT, and incubated for 10 min at 4 °C. Chromatin was isolated by centrifugation at 7,400 rpm for 30 seconds. The pellet was gently resuspended in 1,200 μl of nuclei lysis buffer (50 mM Tris-Cl pH 8.0, 10 mM EDTA, 1 % SDS) with protease inhibitor and 1 mM DTT, incubated for 10 min at 4 °C, and sonicated for 10 cycles (30 sec on/ 30 sec off). The resulting chromatin lysate was centrifuged at 13,000 rpm for 15 min at 4 °C. The supernatant was diluted with 4,800 μl of IP dilution buffer (20 mM Tris-Cl pH 8.0, 2 mM EDTA, 150 mM NaCl, 1 % Triton X-100, 0.01 % SDS), and incubated for 1 h at 4 °C with 60 μg of rabbit IgG, and 60 μl of Protein A magnetic beads for pre-clearing. After pre-clearing, the supernatant was collected for immunoprecipitation.

Immunoprecipitation was performed with 6 ml of pre-cleared chromatin, 200 μl of anti-TEAD4 antibody, and 60 μl of Protein A magnetic beads overnight at 4 °C with rotation. The next day, immunocomplexes were washed sequentially with the following buffers: once with IP Wash I Buffer (20 mM Tris-Cl pH 8.0, 2 mM EDTA, 50 mM NaCl, 1 % Triton X-100, 0.1 % SDS), twice with High salt buffer (20 mM Tris-Cl pH 8.0, 2 mM EDTA, 500 mM NaCl, 1 % Triton X-100, 0.01 % SDS), once with IP Wash II buffer (10 mM Tris-Cl pH 8.0, 1 mM EDTA, 0.25 M LiCl, 1 % NP-40, 1 % Na-deoxycholate), and finally twice with TE pH 8.0. The immunocomplexes were eluted with 200 μl of elution buffer (1 % SDS and 0.1 M NaHCO3) for 30 min at 45 °C with mixing at 1,000 rpm. Cross-links were reversed by adding 0.25 M NaCl and RNase A (1 μg/μl) and incubating overnight at 65 °C. The next day, 4 μl of Proteinase K (Cat# P8107S; NEB) was added and incubated for 2 h at 42 °C. The immunoprecipitated DNA was purified using a QIAquick PCR purification kit (Cat# 28106; QIAGEN) in 50 μl of EB (elution buffer).

### ChIP-seq library construction

ChIP-seq libraries were prepared using the NEXTflexTM ChIP-seq kit (Cat# NOVA-5143-02; PerkinElmer) according to the manufacturer’s instructions. Briefly, 40 μl of purified ChIP DNA was end-repaired, and size-selected (250-300 bp) using AMPure XP beads. The subsequent steps, comprising adenylation, adapter ligation, and PCR amplification, were carried out following the manufacturer’s instructions. Library quality was assessed using a Bioanalyzer (Agilent) with a High Sensitivity DNA chip, showing an average library size of 250–350 bp. For multiplexing, equimolar amounts of libraries were pooled, considering the desired sequencing depth per sample (20–40 million reads per library). ChIP-seq libraries were sequenced on an Illumina NextSeq platform with single-end reads of 76 bp.

### ChIP-seq data processing and analysis

Raw sequencing reads were aligned to the human reference genome (hg19) using Bowtie2 with default parameters, and duplicate mapped reads were removed using SAMtools. TEAD4-binding peaks were first defined in wild-type cells and then compared with YAP/TAZ knockout samples to classify regions as STABLE (peaks showing no significant change in TEAD4 binding), LOSS (peaks with decreased TEAD4 binding in YAP/TAZ knockout cells), or GAIN (peaks with increased TEAD4 binding in YAP/TAZ knockout cells). Heatmaps and metagene plots were generated by calculating read density in 25-bp bins over ±5 kb around TEAD4 peak centers and visualizing the resulting matrices. For genome browser visualization, normalized coverage files (bigWig) were generated from ChIP-seq alignments, and known motif enrichment in TEAD4 LOSS regions was assessed using HOMER with default settings.

### Comet assay

Alkaline comet assays were performed using the OxiSelect™ Comet Assay Kit (Cell Biolabs). Agarose-precoated slides were prepared by dipping glass slides into molten 1% agarose. A total of 2 × 10⁴ cells were washed in ice-cold PBS, resuspended in 250 μL PBS, mixed with 250 μL of 1% low-melting agarose (Invitrogen #16520050), and spread onto the precoated slides. Slides were immersed in lysis buffer (Cell Biolabs) overnight at 4 °C in the dark, then subjected to alkaline electrophoresis at 28 V for 25 min. After air-drying, DNA was stained with 1× GreenView™ DNA Gel Stain and imaged using an EVOS M5000 microscope. Tail moments were quantified using a Python3/SciPy–based analysis pipeline.

### Cell viability assay

Cells were seeded into 12- or 24-well plates and grown to 50–70% confluency. Cells were then treated with the indicated compounds for the durations specified in each figure. Cell viability was assessed by measuring cellular ATP levels using the CellTiter-Glo (Promega, G7570) luminescent assay according to the manufacturer’s instructions, and luminescence was quantified using a microplate spectrometer.

### Colony formation assay

For clonogenic survival under genotoxic stress, 1,000 cells per well were seeded into 24-well plates. After 48h, cells were treated with doxorubicin in the presence or absence of the indicated inhibitors for 7 days. To assess long-term regrowth, cells were maintained for an additional 7 days with medium replacement every 3–4 days. Cells were then fixed with 4% paraformaldehyde (PFA) and stained with 0.05% (w/v) crystal violet for 30 min at room temperature. For well-separated colonies, colony number was quantified. When colonies were confluent or difficult to resolve individually, total colony area was quantified using ImageJ.

### Drug synergy analysis

To evaluate the combinatorial effects of doxorubicin and TEAD inhibitors, 1,000 cells per well were seeded in 96-well plates and treated with the indicated concentrations of each drug, alone or in combination. After 72 h, cell viability was measured using the CellTiter-Glo assay (Promega) according to the manufacturer’s instructions. Drug interaction effects were quantified using the Bliss independence model, which assumes that the effects of two agents are independent and probabilistic. Bliss synergy scores and dose–response interaction maps were computed using SynergyFinder+^87^.

### Hippo pathway transcriptomic and genetic alteration analyses in CCLE

Hippo pathway target genes were defined based on prior literature and included CCN1, CCN2, AMOTL2, ANKRD1, IGFBP3, F3, FJX1, NUAK2, LATS2, GADD45A, TGFB2, PTPN14, NT5E, AXL, DOCK5, ASAP1, RBMS3, MYOF, ARHGEF17, CCDC80, CRIM1, and FOXF2^60^. Transcriptomic features were derived by calculating the mean expression of TEAD1/2/3/4 (Mean TEAD), YAP1 and WWTR1 (TAZ) (Mean YAP/TAZ), and the curated Hippo target gene set for each sample using DepMap expression data (OmicsExpressionTPMLogp1HumanProteinCodingGenes.csv). These metrics were visualized using scatter plots to define global expression relationships and to stratify samples into YAP/TAZ-high and YAP/TAZ-low groups.

Hippo pathway alteration status was defined using mutation and copy number data from DepMap public releases. Damaging mutations were obtained from DepMap Public 25Q3 (OmicsSomaticMutationsMatrixDamaging.csv), and gene-level copy number values were obtained from DepMap Public 24Q4 (OmicsCNGene.csv). A curated set of Hippo pathway genes, including upstream regulators and core kinases (NF2, STK4, STK3, SAV1, MOB1A, MOB1B, LATS1, LATS2, MAP4K1–5, MAP4K4, MINK1, TNIK), was used to define pathway status. Loss-of-function events were defined as damaging mutations, and copy number values ≥ 3 were classified as amplification. Copy number status of YAP1 and WWTR1 (TAZ) was additionally incorporated. Each cell line was classified as Hippo-mutant if at least one gene within the curated set harbored a damaging mutation or amplification, and Hippo–wild-type otherwise.

### Drug sensitivity and Genetic dependency analysis

Drug sensitivity was assessed using area under the dose–response curve (AUC) values from the Genomics of Drug Sensitivity in Cancer (GDSC) datasets (GDSC1 and GDSC2; sanger-dose-response.csv)^59, 88^. Within the YAP/TAZ-low subset, associations between drug response and TEAD expression were evaluated using Spearman correlation between AUC values and Mean TEAD. Volcano plots were generated to visualize correlation coefficients and corresponding adjusted P values across compounds. Differential associations between YAP/TAZ-low and YAP/TAZ-high groups were further assessed by comparing correlation coefficients using Z-score transformation. Genetic dependency was evaluated using gene effect scores from CRISPRGeneEffect.csv (DepMap Public 25Q). Cell lines with gene effect scores < −0.5 were classified as dependent on the corresponding gene.

### TCGA pan-cancer analysis of TEAD expression, genomic instability, and mutation landscape

#### Data acquisition and preprocessing

Genomic and clinical data were obtained from the TCGA Pan-Cancer Atlas (10,967 patients across 33 cancer types). Clinical annotations, including tumor type, aneuploidy score, and fraction of genome altered (FGA), along with batch-normalized mRNA expression data (RSEM), were retrieved for TEAD1–4, YAP1, WWTR1 (TAZ), and curated Hippo pathway target genes. Samples lacking matched clinical, expression, or genomic instability data were excluded.

For each sample, mean expression of gene sets (TEAD1–4, YAP/TAZ, or Hippo target genes) was calculated and log₂-transformed [log₂(mean expression + 1)]. Within each cancer type, patients were stratified into high (≥ median) and low (< median) expression groups to control for inter-tumor heterogeneity.

#### Association with genomic instability

Chromosomal instability was assessed using two TCGA-derived metrics: aneuploidy score and fraction of genome altered (FGA). Differences between high and low expression groups were evaluated independently within each cancer type.

#### Statistical analysis and visualization

Group comparisons were performed using unpaired two-tailed Student’s t-tests for normally distributed data or Mann–Whitney U tests for non-normal distributions. Statistical significance was defined as P < 0.05. Data were visualized as box-and-whisker plots. Data preprocessing and grouping were performed in Python (pandas, numpy), and statistical analysis and visualization were conducted using GraphPad Prism.

#### TEAD alteration frequency and sequence conservation

Somatic mutation data from the TCGA Pan-Cancer Atlas were analyzed to determine alteration frequencies across TEAD paralogs (TEAD1–4). To identify mutation enrichment within conserved functional regions, protein sequences of human TEAD1–4 were aligned, and positional sequence variability was quantified using Shannon entropy. Somatic mutations were mapped onto the aligned sequences, and mutation frequencies were quantified across conserved residues, with particular focus on regions within the TEA DNA binding domain implicated in PAR binding and γH2AX recognition.

#### Immunofluorescence

Cells were seeded onto 18-mm glass coverslips placed in 12-well plates. Coverslips were pre-coated with poly-L-ornithine (1:20 dilution; Sigma P4957) for 15 min at 37 °C, rinsed with PBS, and then used for plating. For immunofluorescence, cells were fixed in 4% paraformaldehyde/PBS for 15 min and permeabilized with 0.2% Triton X-100/PBS for 10 min. To visualize chromatin-associated proteins, a cytoskeleton extraction step was performed prior to fixation. Cells were incubated on ice for 5 min in CSK buffer (10 mM HEPES pH 7.4, 100 mM NaCl, 300 mM sucrose, 1 mM EDTA, 1 mM MgCl₂, 1 mM DTT, 0.2% Triton X-100) supplemented with Halt™ protease and phosphatase inhibitors (Thermo), followed by PBS wash and fixation as above. After permeabilization, cells were blocked in 3% BSA/PBS for 30 min at room temperature and incubated with primary antibodies diluted in 3% BSA/PBS overnight at 4 °C. Following PBS washes, cells were incubated with secondary antibodies for 2 h at room temperature. Nuclei were counterstained with 300 nM DAPI for 5 min, washed with PBS, and mounted using ProLong® Gold antifade reagent (P36930). Images were acquired on a Leica DMI8 microscope using LAS X software.

#### EJ5-GFP assay

NHEJ efficiency was examined sing HEK293A (WT) cells harboring the EJ5–GFP reporter. The EJ5–GFP construct (pimEJ5GFP; Addgene #44026), a gift from Jeremy Stark, was integrated as previously described^55^. To measure loss of function of TEADs in NHEJ, H239A EJ5-GFP cells were transfected with an I-SceI for 48-72h one day after transfection with siTEAD1/2/3/4 or BY116 treatment. The percentage of GFP-positive cells were analyzed using flow cytometry (Thermo, Attune NxT).

#### U2OS 2-6-5 DNA double-strand break reporter cells

U2OS 2-6-5 cells were seeded onto 18-mm glass coverslips and grown to 50–70% confluency. For inhibitor treatments, cells were pretreated for 1 h with ATM inhibitor (KU-60019, 10 μM), ATR inhibitor (VE-821, 10 μM), DNA-PK inhibitor (NU-7441, 5 μM), PARP inhibitor (Olaparib, 10 μM), or 116-10 (10 μM) prior to DSB induction. For transfection experiments, EGFP–TEAD4 WT or mutant constructs were introduced 12 h before DSB induction. Site-specific DNA double-strand breaks were induced by treatment with 4-OHT (1 μM) and Shield-1 (1 μM) for 6 h, followed by fixation and immunofluorescence processing.

#### Laser mircroirradiation

For microirradiation assays, cells were seeded onto 35-mm glass-bottom dishes (Cellvis, D35-20-1.5H). Fluorescently tagged expression constructs were transfected 24 h prior to irradiation. Cells were pre-sensitized with Hoechst (10 µg/mL) for 30 min before laser exposure. Microirradiation was performed on a Leica DMI8 confocal microscope equipped with the FRAP module. Regions of interest (ROIs) were irradiated with a 405-nm UV-A laser at 100% power for 3.6 s, using either a line or circular ROI, and time-lapse images were collected.

#### Chromatin condensation assay

For chromatin condensation measurements, cells were transfected with PAGFP–H2B and subjected to microirradiation as described above. Chromatin expansion was quantified using a Python/Scipy analysis pipeline. Briefly, the PAGFP–H2B–positive region was segmented, and chromatin width was defined as the area of the segmented region divided by the length of the major axis of the minimal bounding rectangle encompassing the PAGFP–H2B signal.

#### Binding site prediction and molecular docking simulations

The binding site of TEAD N-terminal was predicted and generated using the PASSer2.0 protein allosteric sites server with chain A in PDB code 5GZB with the AutoML option^65^. The Glide program in the Schrödinger suite was utilized for BY116 docking to TEAD N-terminal^64^. The protein preparation wizard within the Schrödinger suite was used to prepare the protein (PDB code: 5GZB) for the docking protocol, and the prepared protein structures were used for generating receptor grids. BY116 and BY116-10 were docked using standard precision (SP) mode using Meastro’s Glide.

#### Molecular Dynamics simulations

The MD simulations were performed using Desmond in Schrödinger. The protein-ligand complex resulting from the docking simulation was inserted into an orthorhombic box filled with explicit water molecules (TIP3P model) and a buffer distance of 10 Å. An MD simulation was performed using the OPLS4 force field. Ions (Na+ and Cl–) were added to simulate the physiological concentration of monovalent. ions (0.15 M). NPT (a constant number of particles, pressure, and temperature) uses constant temperature (300 K) and pressure (1.01325 bar) as the ensemble class. The default protocol in Desmond was used to reach system equilibration, and a 2ns simulation was performed for each complex^89^. Trajectory frames from the 200ns simulations (4,000 frames) were subsampled at 10 frame intervals and clustered based on the RMSD of ligand heavy atoms. A representative snapshot was selected from the most populated cluster.

#### Complex structure prediction with AlphaFold3

Complex structures of TEAD4 with γH2AX, poly(ADP-ribose) (PAR), and BY 116 were predicted using Al AlphaFold3 (version 3.0.1). The full-length human TEAD4 (UniProt ID: Q15561) and H2AX (UniProt ID: P16104) sequences were used for all predictions and were provided as separate chains in the AlphaFold3 input. For protein–ligand complex prediction, the chemical structures of PAR and BY 116 were provided in SMILES format. Phosphorylation of γH2AX at Ser140 was explicitly modeled during structure prediction. Detailed definitions of post-translational modifications and ligand chemical inputs are provided in the Supplementary Methods. Structure prediction was performed using multiple independent random seeds. All output models were ranked using the AlphaFold3 internal ranking score, and the top-ranked model was selected for downstream analyses. Model confidence was evaluated using inter-chain predicted TM-score (ipTM), global predicted TM-score (pTM), and predicted aligned error (PAE). Structural visualization and interaction analyses were performed using Schrödinger Maestro and PyMOL. Detailed information regarding the computational environment, containerization, database preparation, multiple sequence alignment generation, inference parameters, and model selection procedures is provided in the Supplementary Methods.

---

**Figure S1.**
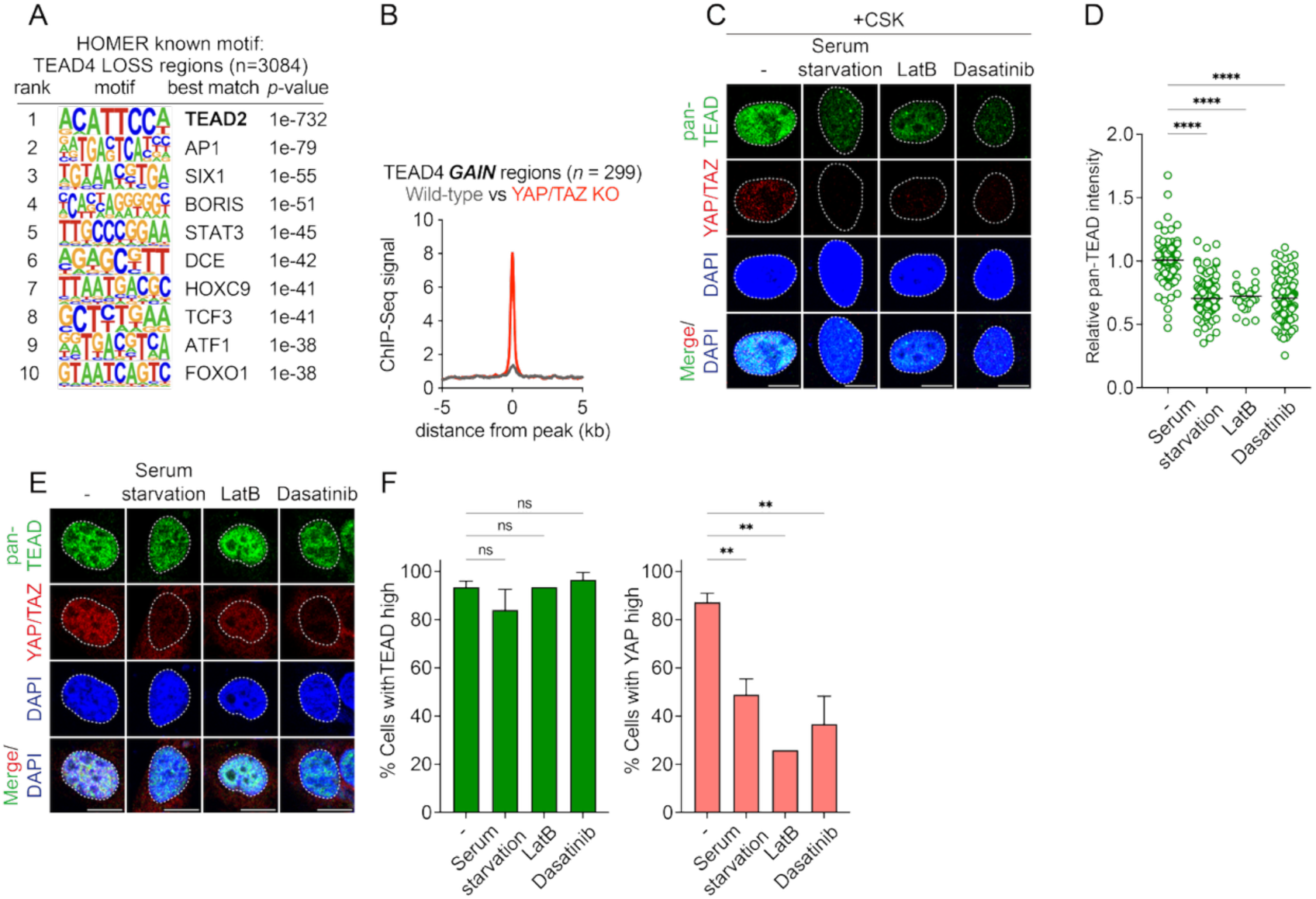
Canonical chromatin engagement of TEAD depends on YAP/TAZ availability. (A) HOMER known-motif enrichment analysis of TEAD4-loss regions (n = 3,084) defined in Figure 1C. The canonical TEAD motif (TEAD2) is the top-ranked hit, confirming the selective loss of TEAD binding at canonical Hippo target sites upon YAP/TAZ knockout. (B) Average TEAD4 ChIP-seq signal density at TEAD-gain regions (n = 299). (C) Confocal images of U2OS cells treated with serum starvation (–serum, 12 h), latrunculin B (0.1 µg/mL, 1 h), or dasatinib (5 µM, 6 h), stained for pan-TEAD (green) and YAP/TAZ (red) after CSK pre-extraction to enrich chromatin-associated proteins. (D) Quantification of nuclear pan-TEAD intensity in (C), normalized to untreated controls (NT: n=75; serum starvation: n=112; LatB: n=26; dasatinib: n=99). Data pooled from three independent experiments. (E) Confocal images of U2OS cells treated as in (C) but without CSK extraction, stained for pan-TEAD (green) and YAP/TAZ (red). (F) Fraction of cells exhibiting nuclear pan-TEAD or YAP/TAZ intensities above one-third of the control mean in (E). Data represent pooled biological replicates (NT/serum starvation/LatB/dasatinib: n=3, 3, 1, 3

**Figure S2.**
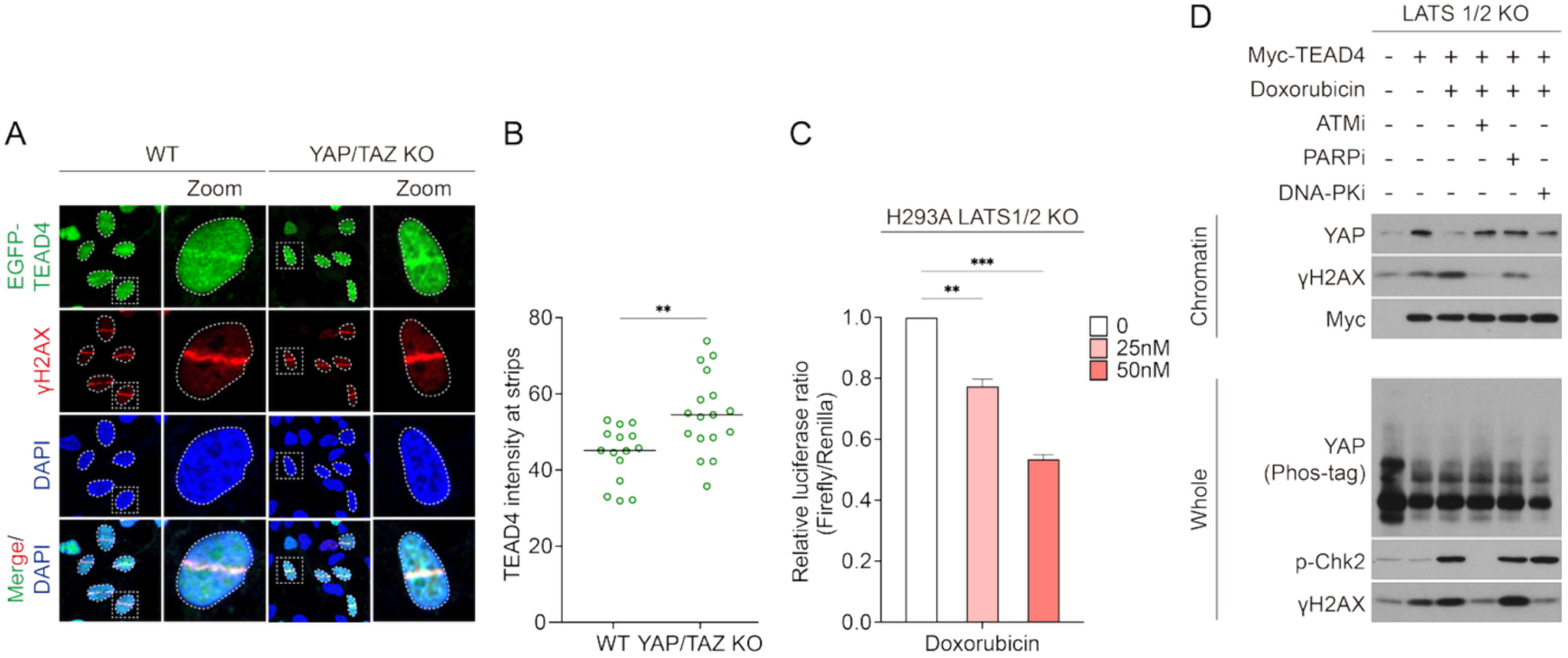
TEAD recruitment to DNA lesions is independent of YAP/TAZ and canonical Hippo signaling. (A) Confocal images of EGFP–TEAD4 (green) and γH2AX (red) in HEK293A wild-type and YAP/TAZ-knockout cells 2 h after microirradiation. (B) Quantification of EGFP–TEAD4 enrichment at laser-induced damage tracks from (B). Lines indicate medians (WT, n = 15; YAP/TAZ-KO, n = 17; unpaired two-tailed t-test). (C) 8×GTIIC luciferase reporter assay measuring YAP/TAZ–TEAD transcriptional activity in HEK293A LATS1/2-knockout cells treated with doxorubicin (25 nM or 50 nM) for 36 h. Data represent two independent biological replicates. (D) Immunoblot analysis of chromatin and whole-cell fractions from HEK293A LATS1/2-knockout cells transfected with Myc–TEAD4 and pretreated with ATM inhibitor (KU-60019, 5 μM), PARP inhibitor (olaparib, 5 μM), or DNA-PK inhibitor (NU-7441, 5 μM) for 30 min prior to doxorubicin treatment (5 μM, 3 h).

**Figure S3.**
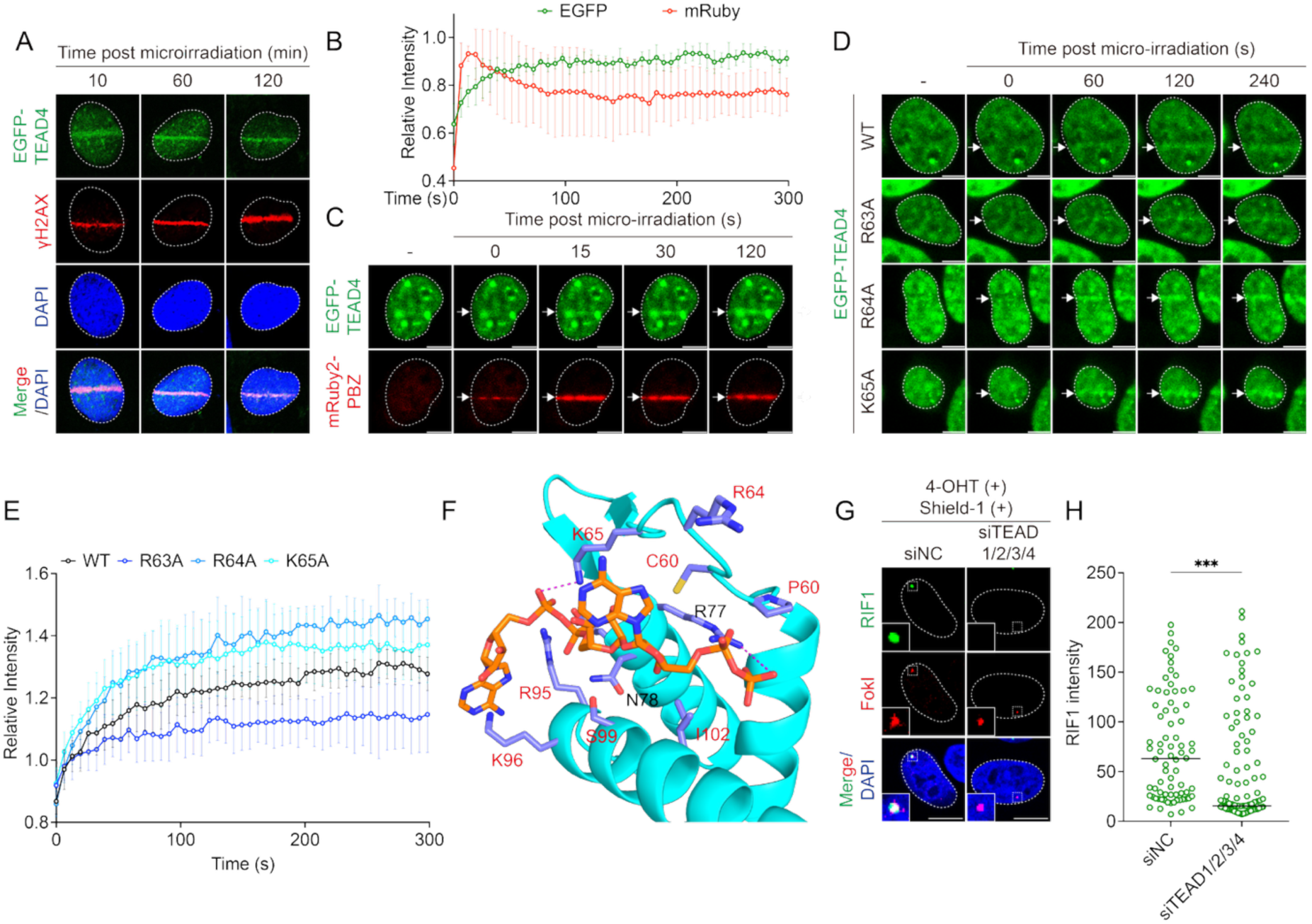
TEAD recruitment to DNA lesions requires the TEA domain and PAR-recognition motifs. (A) Confocal images showing co-localization of EGFP–TEAD4 (green) with γH2AX (red) at laser-induced DNA damage tracks at 10, 20, and 120 min after microirradiation. (B) Quantification of EGFP–TEAD4 and mRuby2–PBZ fluorescence intensities normalized to their respective maximum signals over time (mean ± SD; n = 3). (C) Live-cell imaging showing recruitment dynamics of EGFP–TEAD4 (green) and the PAR sensor mRuby2–PBZ (red) at laser-induced DNA damage sites. White arrows indicate irradiated regions. (D) Live-cell imaging of EGFP–TEAD4 WT, R63A, R64A, and K65A expressed in HEK293A TEAD1/2/4-knockout cells following microirradiation. White arrows indicate irradiated regions. (E) Quantification of recruitment kinetics for the TEAD4 mutants shown in (D), normalized to pre-irradiation intensity (WT, n = 7; R63A, n = 9; R64A, n = 5; K65A, n = 9). (F) AlphaFold3 modeling performed in a Singularity container using a non-canonical ligand representation of PAR. Model confidence metrics were high (ranking_score = 0.96; ipTM = 0.82; pTM = 0.77), with chain-pair ipTM values up to 0.82 and low-to-moderate minimum chain-pair PAE values (1.76–3.33 Å). No steric clashes were detected (has_clash = 0), and the predicted disorder fraction was 0.30. (G) Representative confocal images of U2OS 2-6-5 cells induced with 4-OHT and Shield-1 for 6 h to generate FokI-induced DSBs. Cells were transfected with non-targeting control siRNA (siNC, 20 nM) or pooled siTEAD1/2/3/4 (20 nM total). Accumulation of RIF1 (green) at mCherry–FokI (red) foci is shown. (H) Quantification of RIF1 fluorescence intensity at FokI-positive regions. Horizontal lines indicate medians (siNC, n = 67; siTEAD1-4, n = 47). Unless otherwise indicated, scale bars = 5 μm. Statistical significance was determined by one-way ANOVA: ns, not significant; *P < 0.05; **P < 0.01; ***P < 0.001; ****P < 0.0001.

**Figure S4.**
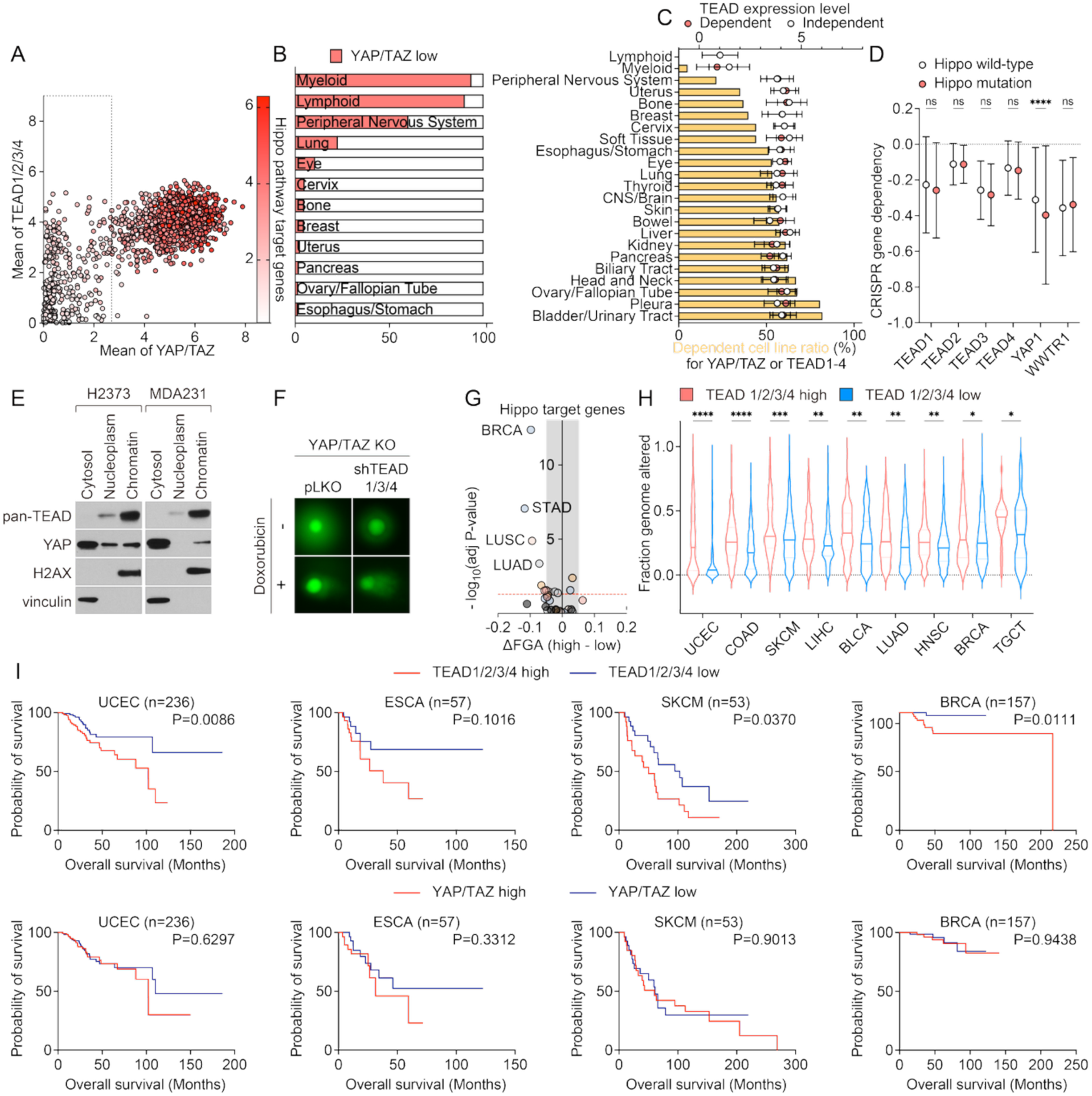
TEAD expression correlates with genomic instability and poor clinical outcome in YAP/TAZ-dispensable cancers. (A) Scatterplot of CCLE cell lines showing the relationship between the mean expression of TEAD1–4 (y-axis) and YAP/TAZ (x-axis). Points are colored using a red gradient representing the mean expression of canonical Hippo pathway target genes. The dashed box indicates YAP/TAZ-low cell lines (YAP/TAZ < 2.5). (B) Distribution of CCLE cancer types within the YAP/TAZ-low group highlighted by the dashed box in (A). (C) Hippo pathway dependency across cancer types. Hippo dependency was defined as the presence of at least one CRISPR gene-effect score < –0.5 for YAP1, WWTR1, TEAD1, TEAD2, TEAD3, or TEAD4. Top: mean TEAD expression in Hippo-dependent (pink) versus Hippo-independent (white) cancers. Bottom: fraction of Hippo-dependent cell lines within each cancer type (yellow bars) (D) CRISPR gene-effect scores for TEAD1–4, YAP1, and WWTR1 in Hippo–wild-type versus Hippo-pathway-mutant cancers. Hippo-pathway mutations were defined as loss-of-function alterations in upstream Hippo regulators or amplification of YAP/TAZ. (E) Cellular fractionation (cytosolic, nuclear, and chromatin fractions) of Hippo-mutant cancer cell lines (H2373 and MDA-MB-231). (F) Representative comet assay images following doxorubicin treatment in YAP/TAZ-knockout cells expressing control vector (pLKO) or shRNAs targeting TEAD1/3/4. (G) Volcano plots showing the association between mean expression of Hippo pathway target genes and fraction of genome altered (FGA) across cancer types. Each point represents a tumor type. The shaded region denotes small effect sizes (|ΔFGA| < 0.05), and the red dashed line indicates adjusted P < 0.05. Significant cancer types are annotated. (H) Fraction of genome altered (FGA) in TCGA tumors stratified by TEAD1–4 (high vs. low, median split). The top nine tumor types showing a significant difference between TEAD-high and TEAD-low groups are shown (UCEC, COAD, SKCM, LIHC, BLCA, LUAD, HNSC, BRCA, TGCT). TEAD or YAP/TAZ-high group n = 255/211/217/178/198/244/251/519/75; TEAD or YAP/TAZ-low group n = 255/210/216/177/197/243/251/518/74. (I) Kaplan–Meier survival analysis of patients treated with radiation (UCEC, ESCA, SKCM) or doxorubicin (BRCA), stratified by TEAD1–4 or YAP/TAZ expression (cutoffs: UCEC = median; ESCA, SKCM, and BRCA = upper versus lower quartiles).

**Figure S5.**
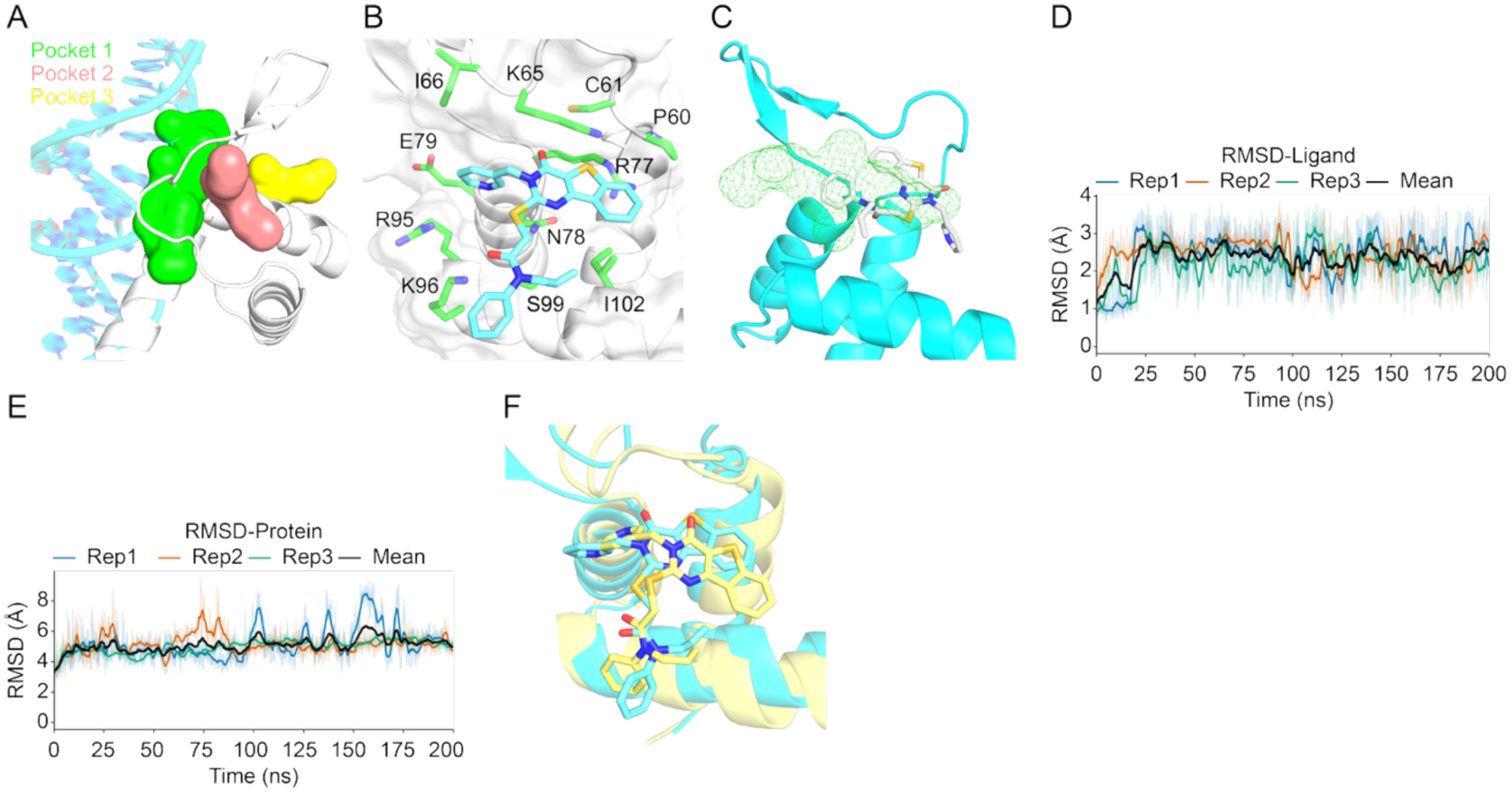
Structural modeling defines an allosteric pocket in the TEAD N-terminus. (A) Visual representation of predicted allosteric binding pockets in the TEAD N-terminal domain. The TEAD N-terminal domain is depicted as a grey cartoon and DNA as an orange helix. Pocket 1 is shown as a green surface, pocket 2 as a yellow surface, and pocket 3 as a blue surface. (B) Predicted binding conformation of compound BY116 within the TEAD N-terminal pocket obtained from molecular docking followed by molecular dynamics (MD) simulation. Amino acid residues within 4 Å of BY116 are shown. (C) Predicted binding pose of compound 116 from AlphaFold3 modeling shown as a white stick representation. The TEAD4 N-terminal domain is shown as a cyan cartoon, and the predicted binding pocket is indicated by a green mesh. (D, E) Root mean square deviation (RMSD) of compound–TEAD complex trajectories during MD simulations. Colored traces indicate three independent replicas (Rep1–Rep3), and the black trace represents the mean RMSD across replicas. Light traces show raw RMSD values, whereas darker traces show smoothed RMSD values. (F) Representative MD simulation snapshot (yellow) shown together with the initial conformation (cyan).

**Figure S6.**
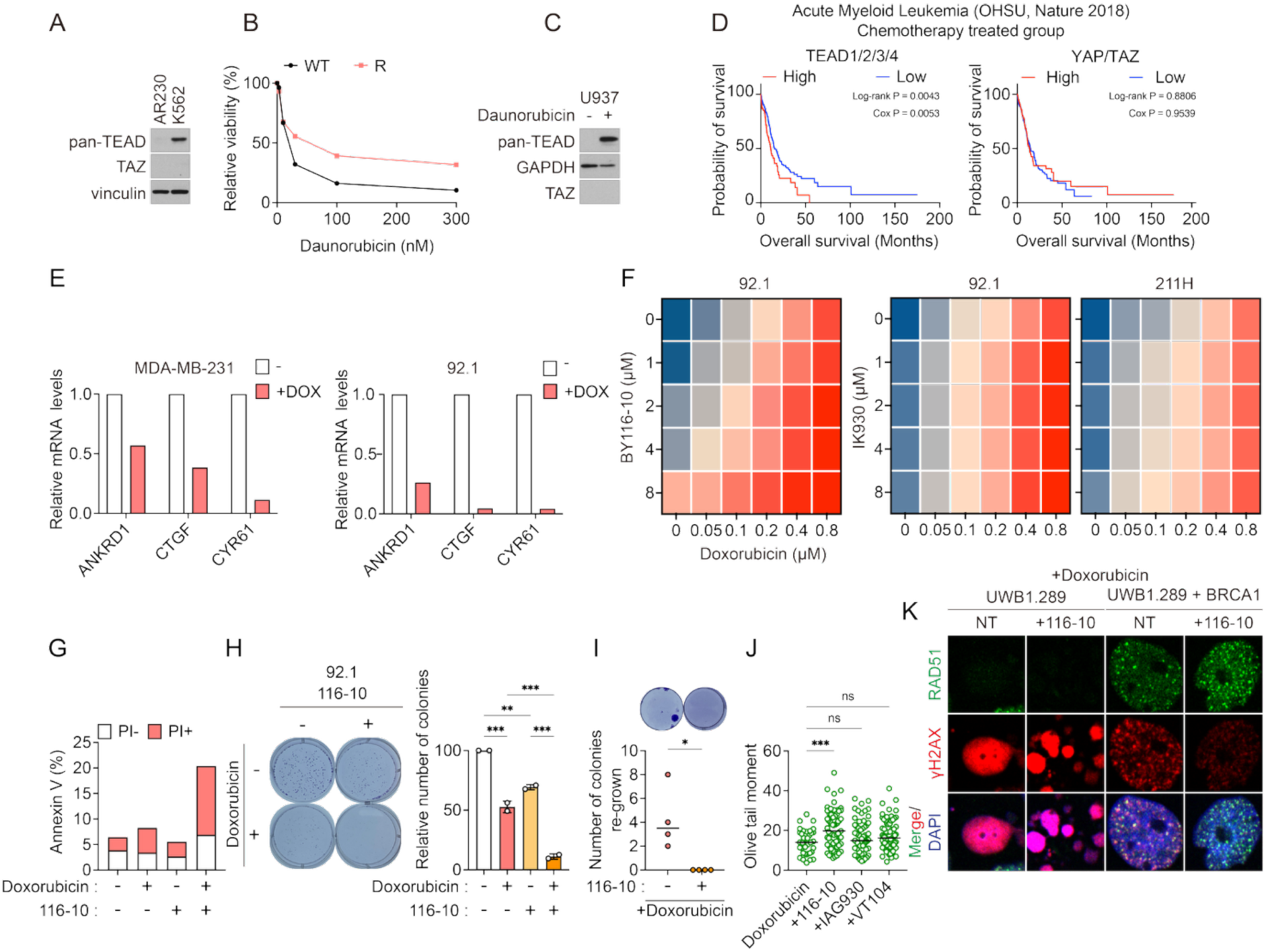
TEAD N-terminal inhibition selectively synergizes with DNA double-strand break–inducing agents. (A) Immunoblot showing endogenous pan-TEAD expression levels in AR230 and K562 cells. (B) Daunorubicin sensitivity in U937 wild-type and daunorubicin-resistant cell lines. (C) Immunoblot showing endogenous pan-TEAD expression levels in U937 wild-type and daunorubicin-resistant cells. (D) Kaplan–Meier survival analysis showing that the mean expression of TEAD1–4 correlates with poor prognosis in AML patients treated with chemotherapy, whereas the mean expression of YAP/TAZ shows no such association. (E) qPCR measurement of Hippo pathway target genes (ANKRD1, CTGF, CYR61) following doxorubicin treatment in MDA-MB-231 and 92.1 cells. (F) Bliss synergy map showing the interaction between doxorubicin and BY116 in 92.1 cells or IK930 in 92.1 and MSTO-211H cells. (G) Annexin V/PI staining showing apoptosis in cells treated with doxorubicin with or without 116-10 in MDA-MB-231 cells. (H) Clonogenic survival assay in 92.1 cells treated with doxorubicin and/or 116-10. (I) Number of colonies re-grown in doxorubicin treatment with addition of 116-10 or not in MDA-MB-231 cells. (J) Olive tail moment measured by comet assay treated doxorubicin with addition of non-treat, 116-10, IAG930, VT104 (all of 1uM) in Hippo-wild-type SUIT2 cells. (K) Immunofluorescence images of UWB1.289 cells (BRCA1-deficient) and BRCA1-restored counterparts treated with doxorubicin or the combination with 116-10, co-stained with RAD51 (green) and γH2AX (red).

